# The Burden of Reliability: How Measurement Noise Limits Brain-Behaviour Predictions

**DOI:** 10.1101/2023.02.09.527898

**Authors:** Martin Gell, Simon B. Eickhoff, Amir Omidvarnia, Vincent Küppers, Kaustubh R. Patil, Theodore D. Satterthwaite, Veronika I. Müller, Robert Langner

## Abstract

Major efforts in human neuroimaging strive to understand individual differences and find biomarkers for clinical applications by predicting behavioural phenotypes from brain imaging data. An essential prerequisite for identifying generalizable and replicable brain-behaviour prediction models is sufficient measurement reliability. However, the selection of prediction targets is predominantly guided by scientific interest or data availability rather than reliability considerations. Here we demonstrate the impact of low phenotypic reliability on out-of-sample prediction performance. Using simulated and empirical data from the Human Connectome Projects, we found that reliability levels common across many phenotypes can markedly limit the ability to link brain and behaviour. Next, using 5000 subjects from the UK Biobank, we show that only highly reliable data can fully benefit from increasing sample sizes from hundreds to thousands of participants. Overall, our findings highlight the importance of measurement reliability for identifying brain–behaviour associations from individual differences.

## Introduction

Major ongoing efforts in human neuroimaging research aim to understand individual differences and identify biomarkers for clinical applications. One particularly promising approach in this regard is the prediction of clinically relevant phenotypes in individuals (e.g. symptoms, treatment response, intellectual abilities) from functional and structural brain measurements (Gabrieli, Ghosh, & Whitfield-Gabrieli, 2015; Woo, Chang, Lindquist, & Wager, 2017; Varoquaux & Poldrack, 2019). Patterns of (resting-state) functional connectivity, the statistical relationship between regional time courses of brain activity (most often expressed as Pearson’s correlation), have been among prominent brain features used for prediction of such phenotypes (Castellanos, Di Martino, Craddock, Mehta, & Milham, 2013; Finn et al., 2015). A large amount of research has focused on the development and improvement of predictive modelling approaches (Shen et al., 2017; Pervaiz, Vidaurre, Woolrich, & Smith, 2020; Kong et al., 2021). Thus far, however, accuracies have remained too low to provide major insights into neural substrates of individual differences or reach clinical relevance (Eickhoff & Langner, 2019; Sui, Jiang, Bustillo, & Calhoun, 2020; Finn, 2021; Tian & Zalesky, 2021; He et al., 2022).

An essential prerequisite for identifying replicable brain-behaviour associations is sufficient reliability of measurements (Vul, Harris, Winkielman, & Pashler, 2009; Milham, Vogelstein, & Xu, 2021). In psychometrics, reliability broadly reflects the consistency of scores across replications of a testing procedure (Standards for Educational and Psychological Testing, 2014). In the context of individual differences, test-retest reliability has received the most attention. It is understood as the degree to which a measure ranks individuals consistently across multiple occasions (i.e. low performers remain low performers on repeated testing). Note that this assumes the measure in question assesses a stable characteristic of the individual or the amount of change between occasions does not differ between individuals (e.g., due to practice from repeated testing). Test-retest reliability is typically evaluated by intraclass correlation (ICC) which is the ratio of between-subject variance and total variance, composed of between-subject, within-subject and error variances (see McGraw and Wong (1996) for a detailed discussion). Measurement noise, understood as the random variability that produces a discrepancy between observed and true values (or repeated observations) is therefore tightly related to reliability as it contributes to error variance in the calculation of ICC. That is, a high level of noise results in low reliability if between-subject variance is held constant. ICC can range from 1 to 0 and is often interpreted as excellent for ICC > 0.8, good for 0.6 – 0.8, moderate for 0.4 – 0.6 and poor for < 0.4 (Landis & Koch, 1977; Hedge, Powell, & Sumner, 2018).

While a large amount of focus has been put on assessing the reliability of brain-based measures (Elliott et al., 2020; Hedges et al., 2022; Noble, Scheinost, & Constable, 2019) and ways to improve them (Finn et al., 2017; Vanderwal et al., 2017; Amico & Goñi, 2018; Li et al., 2019; Pervaiz et al., 2020; Noble, Scheinost, & Constable, 2021), the reliability of behavioural assessments used as prediction targets has been largely neglected. Selecting scientifically or clinically relevant targets for prediction is often guided by pragmatism and logistic constraints (e.g., dataset availability), rather than reliability considerations or criterion validity. Furthermore, classical experimental paradigms may not be well suited for the investigation of individual differences as between-subject variance in such paradigms is often low by design, resulting in low reliability (Hedge et al., 2018). Finally, current assessments of the test-retest reliability of behavioural measures commonly used in the literature show that most fall below the ‘excellent’ reliability (Enkavi et al., 2019; Hedge et al., 2018) that is required for clinical applications (Landis & Koch, 1977; Cicchetti & Sparrow, 1981; Barch & Carter, 2008; Streiner, Norman, & Cairney, 2015). A recent meta-analysis by Enkavi and colleagues (2019) showed the average reliability of 36 tasks assessing self-regulation was on the border between good and moderate (ICC = 0.61), and newly collected data for the same tasks showed even poor reliability (ICC = 0.31). Similarly, assessments of reliability in large datasets and longitudinal samples have reported lower estimates than those reported in test manuals, which often report reliability assessed over relatively short retest intervals (Han & Adolphs, 2020; Taylor et al., 2020; Anokhin et al., 2022).

High measurement reliability is essential as it attenuates relationships between variables. In classical statistics, this is manifested by setting an upper bound on effect size (Spearman, 1910). In the context of machine learning, low reliability can have a profound impact on model performance by lowering the signal-to-noise ratio. Label or target noise (akin to measurement noise) reduces the accuracy of classification algorithms (Frenay & Verleysen, 2014) and increases uncertainty in parameter estimates, training time (Zhu & Wu, 2004) as well as the complexity of a given problem (Garcia, de Carvalho, & Lorena, 2015). Due to inadequate reliability, models may fit variance of no interest (e.g., measurement noise) during training. This results in poor generalisation performance or a failure to learn altogether. Therefore, low out-of-sample prediction accuracy may be a consequence of unreliable targets rather than a weak underlying association. This, in turn, can hamper the assessment of brain-behaviour relationships and strongly undermine efforts directed at biomarker discovery.

Due to effect size attenuation, low reliability also increases the sample sizes necessary to identify effects (Nunnally, 1970; Zuo, Xu, & Milham, 2019). Similarly, targets with higher measurement noise require larger training sets to achieve comparable classification accuracy to less noisy targets (Rolnick, Veit, Belongie, & Shavit, 2018; Wang & Tan, 2018). As a consequence, the estimated strength of brain associations with many behavioural phenotypes will be attenuated and require very large samples to become stable (Marek et al., 2022). These considerations make large datasets for biomarker discovery a necessity rather than an advantage, which in turn poses undesirable logistical, financial and ethical challenges.

The recent availability of large neuroimaging datasets (Van Essen et al., 2013; Sudlow et al., 2015; Volkow et al., 2018), along with advances in machine learning, has led to population-level investigations of brain-behaviour relationships beyond simple correlations. Using a simulation approach and empirical data from four large-scale datasets, we explore how the test-retest reliability of behavioural phenotypes affects their out-of-sample prediction accuracy from functional connectivity and illustrate the tradeoff between reliability and sample size.

## Results

### Low phenotypic reliability reduces the accuracy of brain-behaviour predictions

To systematically test the impact of target reliability on out-of-sample prediction accuracy, we simulated behavioural assessments with reduced test-retest reliability using empirical data from the Human Connectome Project Aging dataset (HCP-A). Reliability was manipulated by incrementally increasing the proportion of random noise within the target variable.

As a proof of principle, we first present results for participant age prediction (n = 647). As expected, systematically reducing the reliability of age resulted in a sharp decrease in accuracy as measurement noise increased (Fig. 1A). Crucially, every 0.2 drop in reliability reduced R^2^ on average by 25%. Mean absolute error (MAE) and correlation of predicted and observed scores followed a similar pattern (Supplementary Results Figure 1). The observed rate of change in accuracy replicated in the UK Biobank dataset (UKB; Supplementary Results Figure 2) and was robust to variations in parcellation (Supplementary Results Figure 3) and algorithm choice (Supplementary Results Figure 4). Reducing the reliability of resting-state functional connectivity by shortening scan duration reduced the overall prediction accuracy, but did not impact the pattern of change in R^2^ (Supplementary Results Figure 5 and 6).

**Figure 1.**
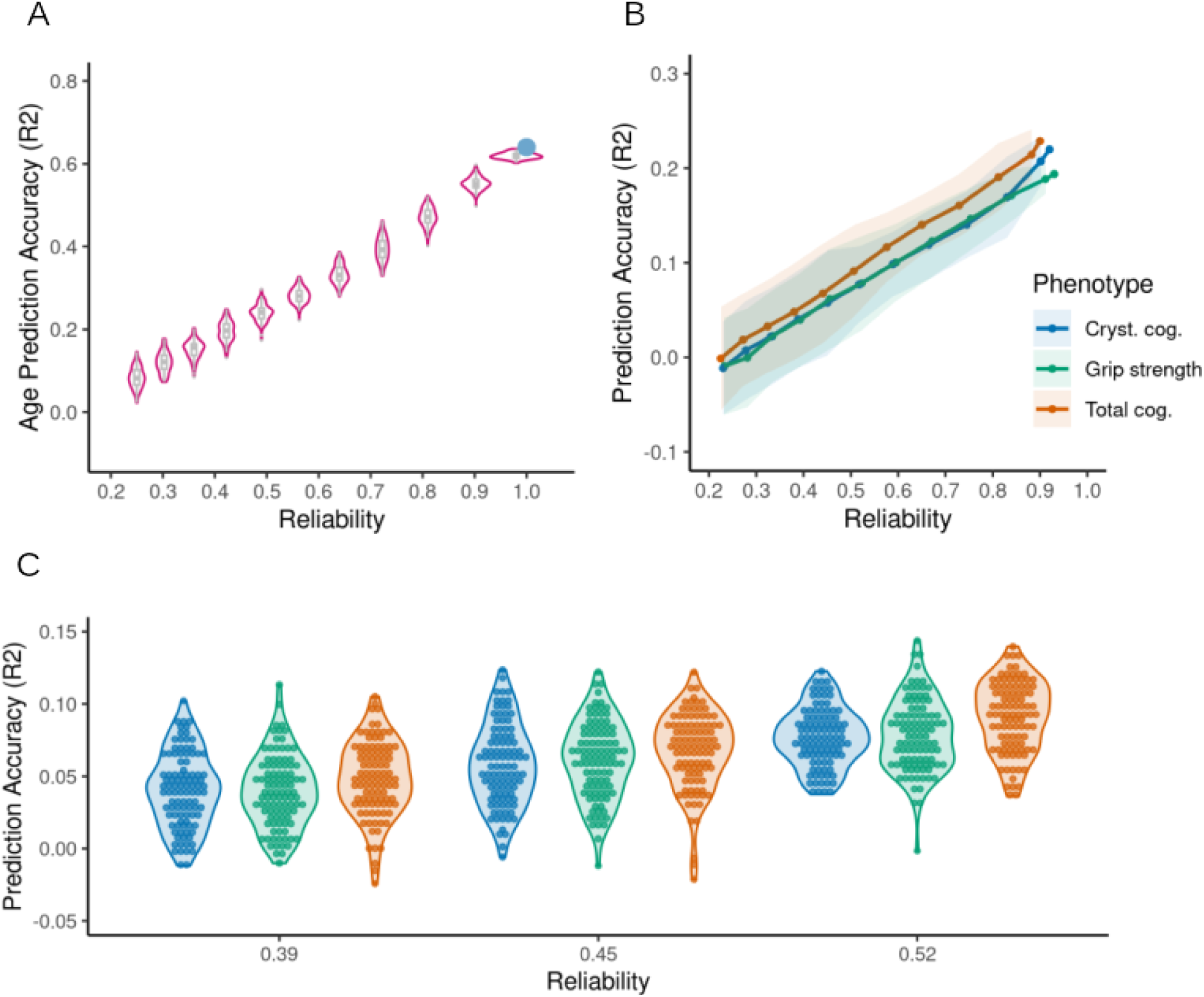
Impact of reliability on prediction accuracy in the HCP-A dataset. (A) Impact of directly reducing the reliability of Age on prediction accuracy (amount of target score variance explained by predicted scores as indicated by R^2^). (B) Impact of reducing the correlation between original and simulated target scores (reflecting reduced reliability) on accuracy in prediction of total cognition composite score, crystallised cognition composite score and grip strength. Solid lines represent the mean across all 100 simulated datasets in each correlation band, shaded areas represent 2 standard deviations in prediction accuracies. (C) Effect of random noise on variability in prediction accuracy. The colour legend is common for panels B and C.

**Figure 2.**
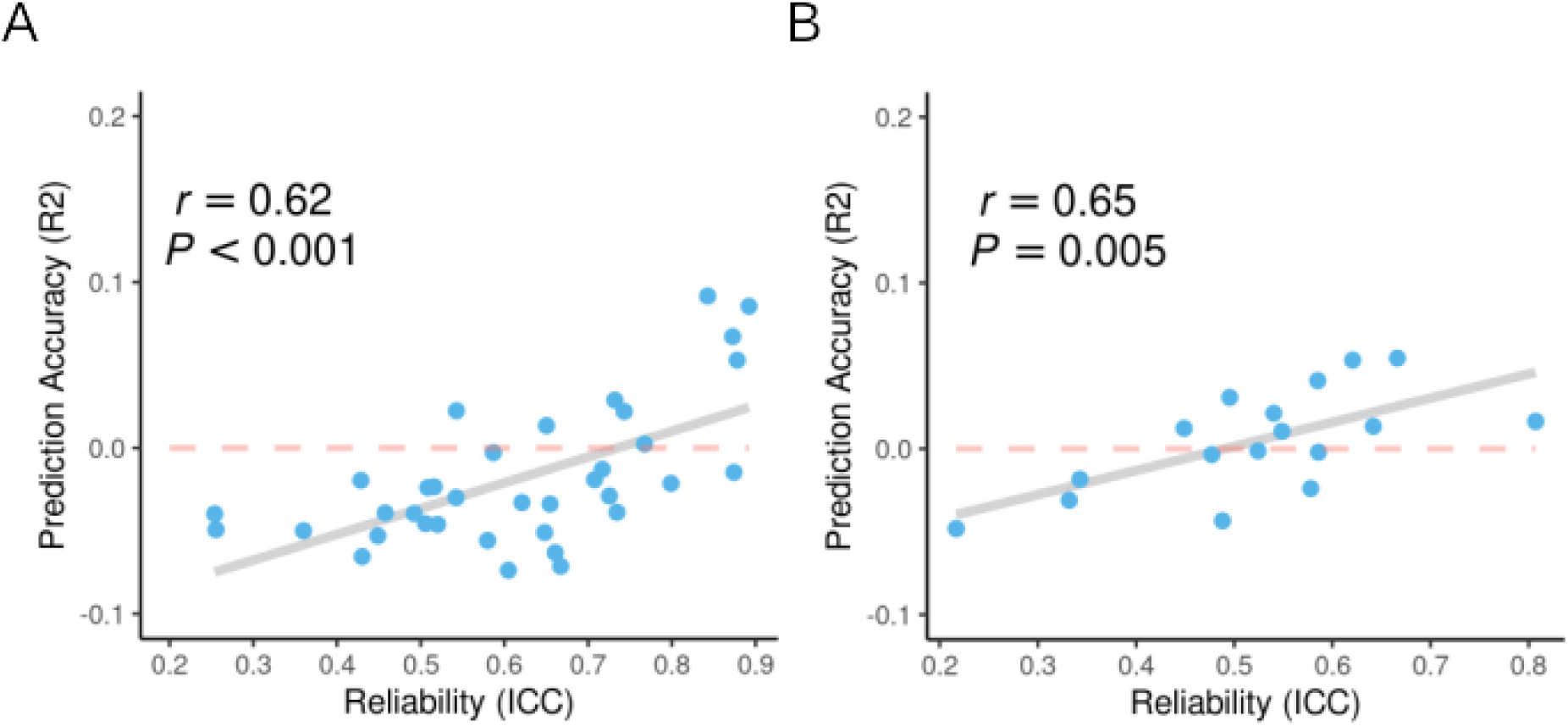
Association between reliability and prediction accuracy. (A) HCP-YA and (B) UKB dataset. Each data point represents a behavioural assessment in each dataset.

**Figure 3.**
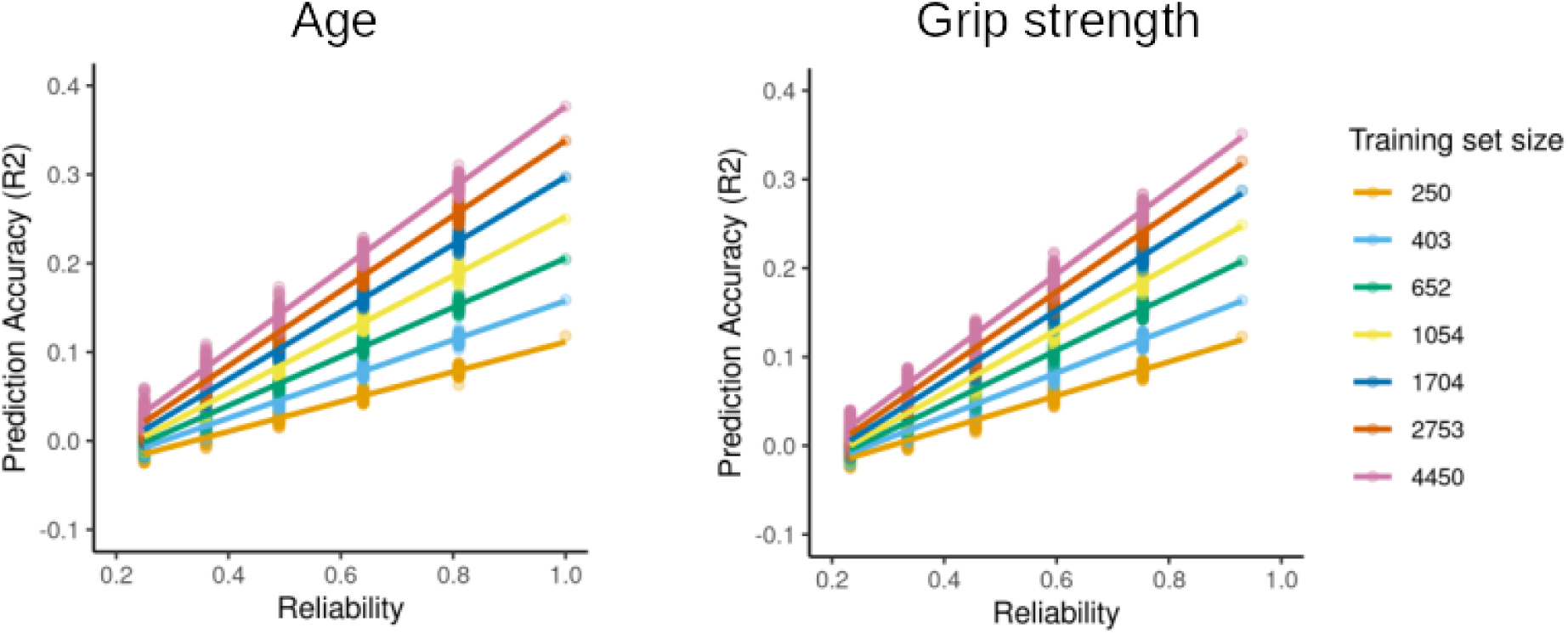
**Prediction and subsampling in UKB**. Impact of training set size on original and simulated data with reduced reliability. Results were fitted with a linear function for illustration purposes.

**Figure 4.**
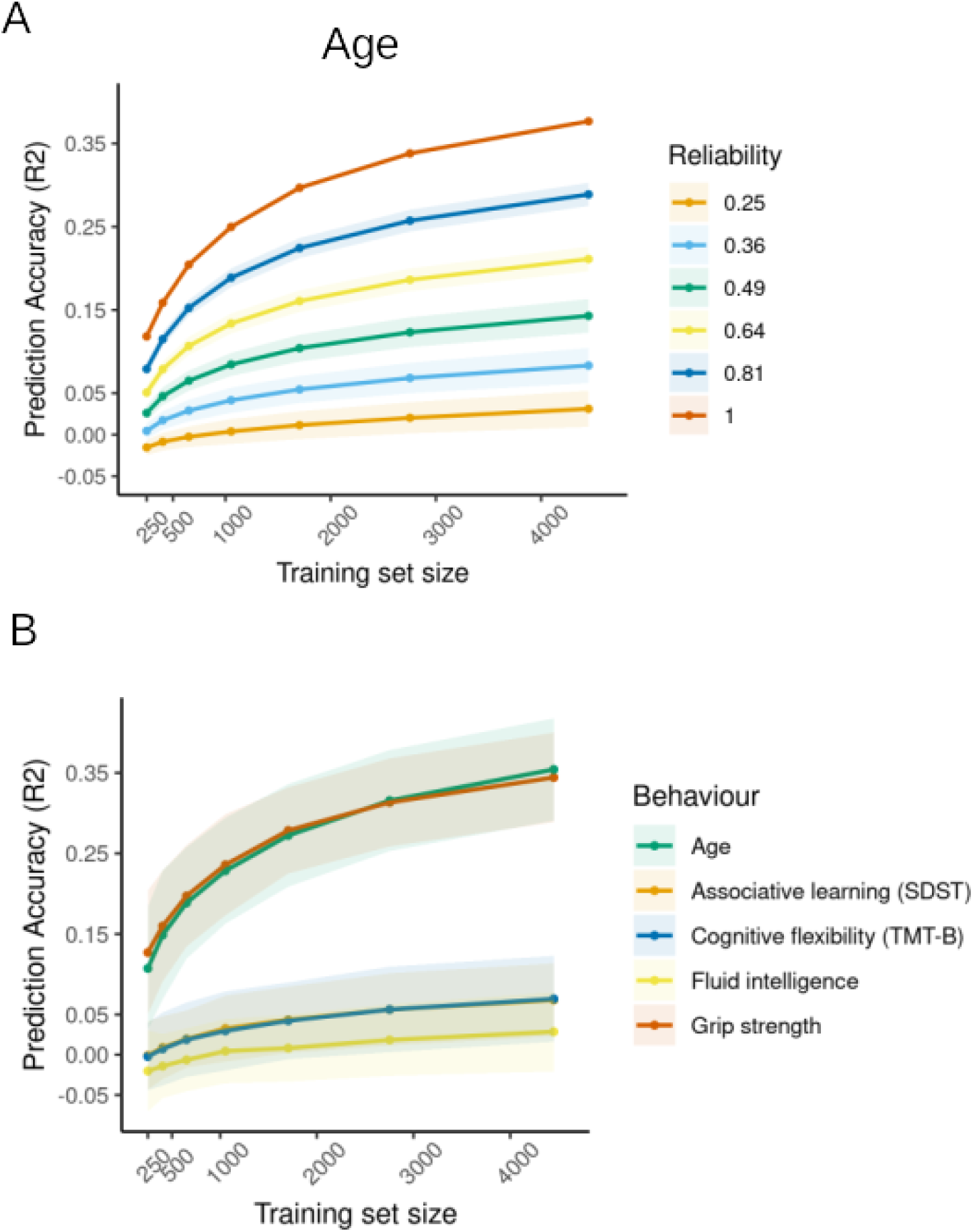
Improvement in prediction accuracy scales with sample size. (A) Impact of training set size on age prediction accuracy in empirical and simulated data with varying levels of reliability. Solid lines represent the mean across all 100 simulated datasets in each correlation band and shaded areas represent 2 standard deviations in prediction accuracies. (B) Impact of training set size on prediction accuracy of empirical behaviours. Solid lines represent the mean accuracy across 100 subsamples and shaded areas represent 2 standard deviations in prediction accuracy. Abbreviations: SDST, Symbol Digit SubstitutionTest; TMT-B, Trail-making task part B.

Next, we investigated the attenuation of prediction accuracy that can be expected in typical studies of brain-behaviour associations, by systematically adding noise to the most reliable measures (ICC ≈ 0.9) available in the HCP-A dataset (n = 550, Supplementary Table 4). This way we simulated new phenotypes with reliabilities that are common in neuropsychological assessments and have plausible true effect sizes. Total cognition could be predicted with an accuracy of R^2^ = 0.23 (MAE = 10.37), crystallised cognition with R^2^ = 0.22 (MAE = 10.24) and grip strength with R^2^ = 0.19 (MAE = 9.79). Similarly to age, reducing their reliability resulted in a decrease in prediction accuracy (Fig. 1B). For all three assessments, R^2^ halved when simulated data reached reliability of 0.6 (R^2^ = 0.12; R^2^ = 0.1; R^2^ = 0.1). Importantly, analysis choices such as confound regression, feature space or feature reliability resulted in small variations in prediction accuracy on empirical and simulated data but had no impact on the rate at which performance decreased (Supplementary Results Figure 7-9). For MAE and correlation between predicted and observed scores see Supplementary Results Figure 10.

We note that prediction accuracy could vary by 0.1 – 0.2 of R^2^ between the best and worst-performing simulated datasets for the same level of noise depending on sampling variability. When reliability reached 0.7 – 0.5, such variability could lead to large fluctuations in accuracy depending on the variable (R^2^ = 0 – 0.12), which in turn could warrant different conclusions regarding the success of predictions (Fig. 1C). All results were corrected by the reliability of phenotypes estimated in previous work (ICC_total cog._ = 0.9 Heaton et al. (2014); ICC_crystalized cog._ = 0.86 Heaton et al. (2014); ICC_grip strength_ = 0.93 Reuben et al. (2013)). As these can vary between studies, we also provide uncorrected results assuming perfect reliability of phenotypes to display more general trends in Supplementary Results Figure 11.

### Target reliability is related to prediction accuracy

Next, we directly investigated the relationship between reliability and brain-behaviour prediction accuracy in empirical data where reliability could be estimated. Using test-retest data from the Human Connectome Project dataset Young Adult (n = 46; HCP-YA) and follow-up data from the UKB dataset (n = 1890), we estimated the reliability of 36 behavioural assessments in HCP-YA (ICCs = 0.25 – 0.89; median ICC = 0.63; Supplementary Results Figure 12) and 17 assessments in UKB (ICCs = 0.22 – 0.81; median ICC = 0.54; Supplementary Results Figure 13). The resulting reliability was then correlated with their prediction accuracy in the full sample (HCP-YA = 771; UKB = 5000) (Fig. 2).

Based on our simulation results we expected increasing attenuation of prediction accuracy as assessment reliability decreased. Confirming this, R^2^ displayed a substantial correlation with test-retest reliability in the HCP-YA dataset (r = 0.62, p < 0.001) and the UKB dataset (r = 0.65, p = 0.005) even though retest intervals were longer (mean retest = 2 years and 6 months compared to 5 months in HCP-YA) and also generalised to the ABCD dataset (r = 0.86; p < 0.001; Supplementary Results Figure 14). Given the small number of retest subjects (n = 46) in HCP-YA, we also correlated R^2^ with the lower and upper bounds of the ICC and observed the same relationship (r = 0.61, p < 0.001 and r = 0.54, p < 0.001, respectively). As models with negative R^2^ values may not be comparable in accuracy, we also correlated only models with positive R^2^ with reliability in HCP-YA and found an even stronger correlation (r = 0.71, p = 0.03). Similar to our main analysis, all variables with reliability lower than < 0.6 displayed very low accuracy (R^2^ < 0.02). Conversely, only variables with excellent reliability (the picture vocabulary task, total cognition, grip strength, reading English and crystallised cognition) could achieve R^2^ > 0.05.

### Influence of phenotype reliability on prediction accuracy scales with sample size

Finally, we sought to investigate how the interaction between reliability and sample size impacts brain behaviour prediction. Using 5000 subjects from the UKB dataset we repeated the same simulation approach in geometrically spaced training set sizes ranging from 250 to 4450 subjects. Systematically increasing random noise in age and grip strength resulted in reduced accuracy for all training set sizes and followed the same pattern of R^2^ halving for every 0.4 drop in reliability observed in our previous analysis (Fig. 3). Importantly, a change of 0.2 in reliability had a larger impact on prediction performance than a change in training set size. For age prediction, even samples of 652 subjects with excellent reliability (r = 0.81; R^2^ = 0.15, R^2^ = 0.005) produced comparable accuracy to the full sample (n = 4450) with a moderate level of phenotypic reliability that is common across behavioural assessments (r = 0.49; R^2^ = 0.14, R^2^ = 0.01). For phenotypes displaying weaker association with functional connectivity, this effect was less pronounced, requiring reliable training sets with half the size of the full sample with less reliable data to achieve comparable accuracy (Supplementary Results Figure 15).

Increasing sample size always resulted in higher prediction accuracy irrespective of reliability (Fig. 4). However, the largest improvements in accuracy were observed for highly reliable data, while data with moderate reliability showed only minor gains (Fig. 4A). This was particularly pronounced for samples below 1000 participants. Next, we investigated how prediction accuracy of empirical data with varying levels of reliability increases as a function of training set size (Fig. 4B). Replicating results from simulated data with reduced reliability, phenotypes with excellent reliability (ICC_grip strength_ = 0.81; ICC_age_ ≈ 1.0) displayed a steeper and larger improvement in accuracy as sample size increased. Phenotypes with good reliability ^(ICC^_Associative learning_ ^= 0.62; ICC^_Cognitive flexibility_ ^= 0.67; ICC^_Fluid intelligence_ ^= 0.64)^ showed only minor changes in accuracy with proportionally smaller improvements (Supplementary Results Table 1). These remained unchanged when the maximum training set size was increased by an additional 2500 subjects (Supplementary Results Figure 16).

**Table 1.** Overview of datasets and samples used in main analyses.

| Dataset | Analysis | Sample (Female) | Age Range |
| --- | --- | --- | --- |
| HCP-A | Prediction of simulated data | 647 (351) | 36-89 |
| HCP-YA | Prediction of all phenotypes | 771 (358) | 22-35 |
|  | Test-retest | 46 (32) | 22-35 |
| UKB | Prediction of simulated data and all phenotypes | 5000 (2714) | 48-82 |
|  | Test-retest | 1890 (1012) | 48-79 |
| ABCD | Prediction of simulated data and all phenotypes | 4133 (2123) | 9-10 |
|  | Test-retest | 2102 (1026) | 9-10 |

HCP-A, Human Connectome Project Aging; HCP-YA, Human Connectome Project dataset Young Adult; UKB, UK Biobank.

## Discussion

Here we demonstrate the burden of low phenotypic test-retest reliability on brain-based out-of-sample prediction performance. Our results suggest that especially when associations between brain features and behavioural assessments are weak to moderate, levels of reliability that are common for behavioural phenotypes can substantially attenuate large portions of shared variance. Importantly, this attenuation holds irrespective of feature definition, prediction algorithm or dataset, suggesting that analytical choices have little impact. Furthermore, we show that while a larger sample size increases the accuracy of brain-behaviour predictions, highly reliable data in smaller samples can produce comparable results to large amounts of moderately reliable data, and depending on the size of the true relationship can even outperform it. Following on from these findings, we show that only highly reliable data can fully benefit from increasing sample sizes from hundreds to thousands of participants.

### Phenotypic reliability is important for robust results

The attenuation of a correlation between two variables by their reliability was already described by Charles Spearman in 1910. Here we aimed to demonstrate that machine learning approaches used to identify brain-behaviour associations also suffer from low phenotypic reliability and show its impact on out-of-sample prediction accuracy. Generally, we found that reliability attenuated out-of-sample prediction accuracy in a similar manner to what has been described for correlation (Nunnally, 1970; Vul et al., 2009; Zuo et al., 2019) and classification (Frenay & Verleysen, 2014; McNamara, Zisser, Beevers, & Shumake, 2022).

Building on arguments emphasising the importance of reliability in biomarker research (e.g. Milham et al., 2021), we illustrate the amount of attenuation that can be expected by the reliability of routinely collected neuropsychological assessments available in many large datasets. Our results suggest that moderate reliability (ICC = 0.6 – 0.4) can produce serious attenuations of prediction accuracy irrespective of the dataset, rs-fMRI reliability and analytical choices. Moreover, even good levels of reliability (ICC = 0.6 – 0.8) were found to substantially attenuate brain-behaviour associations. In particular, reducing reliability to ICC = 0.6 diminished prediction accuracy on average by half. Strong relationships (e.g. age) were equally susceptible to strong attenuation but unlike weaker ones could still be predicted with poor reliability. However, current estimates indicate that such large effect sizes for brain-phenotype associations are the exception rather than the rule (Button et al., 2013; Marek et al., 2022). Overall, these results indicate that high test-retest reliability of behavioural phenotypes is crucial to fairly evaluate the potential of neuroimaging for the prediction of individual differences in behaviour.

Supporting previous literature (Hedge et al., 2018; Scott, Sorrell, & Benitez, 2019; Enkavi et al., 2019; Taylor et al., 2020; Fawns-Ritchie & Deary, 2020; Anokhin et al., 2022), most behavioural assessments in the datasets used here (HCP-YA, UKB and ABCD) showed reliabilities within the good to moderate range (median ICC = 0.51; Supplementary Results Figure 17) that were susceptible to high attenuations, despite desirable levels for clinical applications (Barch & Carter, 2008; Streiner et al., 2015). As many large neuroimaging datasets utilise similar measurement instruments (e.g. NIH Toolbox; Weintraub et al., 2013), low prediction accuracies observed in many recent reports may be partly driven by suboptimal reliability of prediction targets (Dubois, Galdi, Han, Paul, & Adolphs, 2018; Li et al., 2019; Pervaiz et al., 2020; Mansour, Tian, Yeo, Cropley, & Zalesky, 2021; Wu et al., 2021; McCormick, Arnemann, Ito, Hanson, & Cole, 2022; Heckner et al., 2023). This in turn limits further insights into interindividual differences in brain function and the search for neuroimaging-based biomarkers. Importantly, our results also suggest that the field can benefit significantly from improving measurement practices and optimising behavioural reliability to increase SNR for predictive modelling and increasing association effect sizes.

The final attenuation of brain-behaviour relationships will be determined by the joint reliability of both neuroimaging features and behavioural targets (Nunnally, 1970; Nikolaidis et al., 2022). Reliability of functional connectivity depends on the network (Tozzi, Fleming, Taylor, Raterink, & Williams, 2020), preprocessing steps (Noble et al., 2019) and scan duration, with longer acquisition leading to greater reliability (Noble et al., 2017, 2019; Cho, Korchmaros, Vogelstein, Milham, & Xu, 2021). The marked difference in age prediction accuracy between HCP and UKB datasets we observed here, is therefore, likely related to differences in rsfMRI acquisition (6 minutes in UKB compared to 26 minutes in HCP-A), in addition to lower precision in reported age in the UKB (measured in years compared to months in HCP). In other words, low phenotypic reliability that produced serious attenuation in the HCP-A dataset is likely to display even greater attenuation in datasets with less reliable fMRI measurements. Therefore, the results shown here may represent an optimistic scenario for the field, as 26 minutes of resting-state images collected over two days is, especially in clinical settings, uncommon. However, we also emphasise that the impact of low phenotypic reliability generalised across datasets as well as when feature reliability was directly manipulated. Therefore, even with exceptionally reliable fMRI measurements, unreliable phenotypes are still likely to substantially attenuate out-of-sample prediction accuracy, as consistent ranking across individuals will be impaired.

In addition to overall low prediction performance for data with less than good reliabilities, we observed a large variance in prediction accuracy in simulated data. Specifically, datasets with moderate and poor reliability showed accuracies that could result in opposite conclusions. For example, at ICC = 0.45, the highest accuracies (R^2^ ≈ 0.1) were comparable to those reported for many behavioural assessments (Sasse et al., 2022), while the worst observed accuracy represented a failure of prediction (i.e., R^2^ < 0). As in our simulations, measurement noise was randomly distributed; these results suggest that even phenotypes with moderate reliability may contain enough noise to produce results that will not replicate. Conversely, the higher the reliability, the lower the risk of the variance in results caused by random noise to reach R^2^ = 0. Our findings, therefore, reinforce the necessity for authors to replicate their prediction results across datasets or validate their models in truly independent samples (Poldrack, Huckins, & Varoquaux, 2020).

### Large samples are necessary but not sufficient

In a recent study, Marek and colleagues (2022) have suggested that investigating brain-phenotype associations requires sample sizes of n > 2000, as sampling variability in small effects can result in imprecise effect size estimates. While cognitive ability and total psychopathology used by the authors as exemplary phenotypes have been reported to have excellent reliability (ICC > 0.9; however, see Tiego et al. (2022) for a discussion), the remaining phenotypes that were assessed have more modest reliabilities (ICC = 0.31 – 0.82) (Han & Adolphs, 2020; Taylor et al., 2020; Fawns-Ritchie & Deary, 2020; Fox, Manly, Slotkin, Devin Peipert, & Gershon, 2021; Anokhin et al., 2022). Given this large variation in reliability, the reported sample size requirement is likely not a one-size-fits-all recommendation (Rosenberg & Finn, 2022), as increasing the reliability of many collected behavioural measurements will result in larger effect sizes, effectively reducing the sample size requirement. Here we demonstrate that depending on the true association strength, highly reliable phenotypes can reach comparable prediction accuracy using samples in the hundreds rather than thousands, as they are less subject to marked attenuation by low reliability. These results suggest that collecting more reliable data may be particularly important for research questions where, assuming cross-sectional design is appropriate, many thousands of participants are difficult to acquire (e.g., specific conditions) and discuss ways to implement this below. However, more importantly, we demonstrate that only reliable phenotypes can fully benefit from improvements in prediction accuracy as training set sizes increase from hundreds to thousands of participants that has been observed previously (Nieuwenhuis et al., 2012; Jollans et al., 2019; Traut et al., 2022). Conversely, measurements with poor reliability are likely suboptimal candidates for big data initiatives as collecting thousands of participants will only yield minor increases in accuracy before saturating. Therefore, improving measurement reliability of appropriately selected phenotypes for associations with neuroimaging features will likely boost predictive (and statistical) power in large datasets. Finally, we note that our findings should not be taken to justify the use of small n studies under the guise of high measurement quality. As long as true associations between behavioural phenotypes and neuroimaging display small effect sizes, very large samples will be necessary. Thus, it is important that on top of considering measurement reliability, researchers continue to follow guidelines for generalisable (Paus, 2010) and reproducible predictive modelling (Varoquaux, 2018; Janssen, Mourão-Miranda, & Schnack, 2018; Scheinost et al., 2019; Poldrack et al., 2020).

Across a broad range of tested variables, empirical reliability (estimated from the datasets) was rarely excellent (5 out of 36 tested in HCP-YA, 0 out of 17 in UKB), replicating observations from other large datasets (Anokhin et al., 2022). Furthermore, empirical reliability was generally lower than that reported at test development (Akshoomoff et al., 2013; Reuben et al., 2013; Weintraub et al., 2013; Heaton et al., 2014). Similar differences in reliability between datasets are not uncommon and may be due to differences in retest intervals (Scott et al., 2019; Taylor et al., 2020; Han & Adolphs, 2020; Anokhin et al., 2022). However, assessments of behaviour in large datasets in particular may be subject to other sources of measurement noise resulting from specifics of big data collection such as site differences, coordinator training, relatively low number of trials designed to lower the burden on participants or shortened versions of validated assessments, and participant fatigue from lengthy acquisition protocols. At the same time, best practices in assessing test-retest reliability during test development are not always adhered to, likely producing further discrepancies between studies (Polit, 2014). We note that the actual test-retest reliability of many measures in large datasets is currently hard to assess as outside of HCP-YA none of the most commonly used datasets like HCP-A, UKB and ABCD have test-retest samples. The inability to assess phenotype reliability in these datasets precludes the possibility to disentangle whether poor model performance is due to measurement error or reflects low effect size. If phenotype reliability is indeed substantially lower in large datasets than that reported at test development, many available datasets may be of limited use for individual-differences research and further increasing sample size (e.g. to biobank levels) will be of little benefit. We therefore urge that moving forward, any attempts at identifying biomarkers must involve careful consideration and thorough assessment of the reliability of behavioural as well as neuroimaging measurements e.g. in re-test samples before data is collected at larger scales and evaluated for predictive power.

### Improving phenotypic reliability

A wealth of literature discusses ways of improving measurement reliability. Prior to acquisition, this can be achieved by opting for a deeper phenotyping design (Gratton, Nelson, & Gordon, 2022) either in the laboratory or by means of ecological momentary assessment (Moskowitz & Young, 2006), introducing more rigorous testing strategies such as collecting more trials (for an overview see Zorowitz & Niv, 2022), taking measures to increase between-subject variance (Xu et al., 2023) or acquiring multiple assessments for data aggregation (Nikolaidis et al., 2022). In already acquired data, researchers should select relevant measurements with the best psychometric properties. For behavioural phenotypes, data reduction techniques such as principal component analysis or summary scores, assuming that error variance and loading of all items on a latent dimension are equal (McNeish & Wolf, 2020), can increase reliability and lead to larger effect sizes than individual items (Lohmann et al., 2021; Tian & Zalesky, 2021; Marek et al., 2022; Ooi et al., 2022; Sasse et al., 2022). Supporting this, composite scores of the NIH toolbox tasks in the HCP datasets were more reliable than individual assessments and reached higher prediction accuracy. Similarly, averaging left and right hand grip strength in the UKB dataset compared to each hand separately lead to improvement in both reliability and accuracy. Comparable increases were also achieved when grip strength was averaged across testing occasions (Supplementary Results Figure 18). If equal item loading on a latent dimension cannot be assumed, reliability can be increased using latent modelling frameworks that account for systematic and unsystematic errors. However, more work is necessary to identify the most cost-effective strategies for optimising the reliability of both measurements without sacrificing measurement validity to increase prediction accuracy. To this end, future research should focus on ways to improve the reliability of already acquired data and evaluate best practices to preserve reliability when acquiring new data at large scales.

Although high reliability of either measurement is necessary for meaningful investigations of prediction accuracy, it is not sufficient. For instance, highly reliable phenotypes that don’t capture a valid representation within the brain are not likely to improve effect sizes. Moreover, many behavioural measurements are validated against other established psychological scales or with specific populations in mind, rather than developed in light of their biological relevance. As a result, they may not be well-suited for investigations of brain behaviour associations, and thus, enhancing their reliability may bring little improvement in effect size. Similarly, structural MRI metrics that display better reliability than functional connectivity (Reuter, Schmansky, Rosas, & Fischl, 2012; Masouleh et al., 2020), are often poorer predictors of many psychological constructs (Ooi et al., 2022) that may instead rely on intrinsic fluctuations in neural activity (Waschke, Kloosterman, Obleser, & Garrett, 2021). Therefore, while optimising measurement reliability offers one possible avenue for improving the investigation of individual differences; it will not guarantee larger effect sizes (Finn & Rosenberg, 2021) or better prediction accuracy, especially if the selection of appropriate phenotypes is neglected.

## Conclusion

The recent availability of large-scale neuroimaging datasets, combined with advances in machine learning, has enabled the investigation of population-level brain-behaviour associations. In this study, we demonstrate that common levels of reliability across many behavioural phenotypes in such datasets can strongly attenuate or even conceal actual associations. This, in turn, can lead to scientifically questionable conclusions about the predictive potential of neuroimaging and hinders clinical translation. Therefore, greater emphasis needs to be placed on refining behavioural phenotyping in large datasets on top of similar efforts directed at neuroimaging. Together, more reliable neurobiological measurements and “markers” of behaviour will be necessary to fully exploit the benefits of big data initiatives in neuroscience, promote the identification of potential biomarkers, and contribute to reproducible science.

## Methods

To investigate the impact of phenotypic reliability on brain-behaviour associations, we used functional connectivity to predict empirical and simulated data with varying levels of reliability. Reliability was manipulated by increasing the proportion of random noise (representing error variance) in our prediction targets. Noise simulations were done using data of the Human Connectome Project Aging dataset (HCP-A) due to its favourable ratio between imaging data quality and variance in phenotypic data with high reliability. As increasing noise for the purposes of our analyses may only be meaningful in highly reliable phenotypes, we selected prediction targets based on their published estimates of reliability: age (ICC ≈ 1.0), grip strength (ICC = 0.93; Reuben et al. (2013)), total cognition composite (ICC = 0.86 – 0.95; Akshoomoff et al. (2013); Heaton et al. (2014)) and crystallised cognition composite (ICC = 0.9; Akshoomoff et al. (2013); Heaton et al. (2014)). Next, three datasets – The Human Connectome Project dataset Young Adult (HCP-YA), UK Biobank (UKB) and Adolescent Brain Cognitive Development (ABCD) were used to investigate the association between reliability and prediction accuracy as test-retest or follow-up behavioural data was available in all datasets (and not in HCP-A). The ABCD was used to investigate if the association between reliability and prediction accuracy generalised to a dataset with a different population and preprocessing steps. Lastly, the UKB sample was used to investigate the interaction between reliability and sample size given the large number of subjects available. To create simulated data, the noise was only manipulated on the most reliable phenotypes available in the dataset: age (ICC ≈ 1.0) and grip strength (ICC = 0.93 – 0.96; Bohanon et al. (2011); Hamilton et al. (1994)). Unfortunately, none of the cognitive assessments exhibited reliability values that were high enough for our purpose, with the highest reliability at r = 0.78 for the trail-making B task (Fawns-Ritchie & Deary, 2020). For an overview of datasets see Table 1.

## Datasets

### Human Connectome Project Aging dataset

For our primary simulation analysis, we used data from the Human Connectome Project Aging dataset (Bookheimer et al., 2019; Harms et al., 2018), obtained from unrelated healthy adults. Only subjects with all four complete runs of resting-state fMRI (rs-fMRI) scans and no excessive head movement (framewise displacement < 0.25 mm, which corresponded to 3SD above the mean) were analysed, resulting in a sample of 647 subjects for age prediction (351 female, ages = 36-89) of approximately 550 had all phenotypic data of interest available (see supplementary methods for exact n for each phenotype).

The HCP scanning protocol involved high-resolution T1w MRI images that were acquired on a 3T Siemens Prisma with a 32-channel head coil using a 3D multi-echo MPRAGE sequence (TR = 2500 ms, 0.8 mm isotropic voxels). The rs-fMRI images were acquired using a 2D multiband gradient-echo echo-planar imaging (TR = 800 ms, 2 mm isotropic voxels). Four rs-fMRI sessions with 488 volumes each (6 min and 41 s) were acquired on two consecutive days, with one anterior-to-posterior and one posterior-to-anterior encoding direction acquired on each day.

### Human Connectome Young Adult dataset

To investigate the relationship between reliability and prediction accuracy we used data from the Human Connectome Project Young Adult dataset (Van Essen et al., 2013), partly consisting of related healthy subjects. Only subjects with all four complete runs of rs-fMRI, no excessive head movement (framewise displacement < 0.3 mm, which corresponded to a displacement of 3SD above the mean) and all phenotypes of interest were included (n = 713, 358 female, ages = 22-35). In total, 36 behavioural phenotypes that were available for all subjects and did not display strong ceiling effects were selected for prediction (see supplementary methods for a full list of phenotypes and their distributions). Standardised scores were used when available. Additionally, a test-retest dataset for subjects with all 36 assessments (n = 46, 32 female, ages = 22-35) was used to estimate phenotypic reliability.

The HCP scanning protocol involved high-resolution T1w MRI images that were acquired on a 32-channel head coil on a 3T Siemens “Connectome Skyra” scanner using a 3D single-echo MPRAGE sequence (TR = 2400 ms, 0.7 mm isotropic voxels). The resting state fMRI images were acquired using whole-brain multiband gradient-echo echo-planar imaging (TR = 720 ms, 2 mm isotropic voxels). Four rs-fMRI sessions with 1,200 volumes each (14 min and 24 s) were acquired on two consecutive days, with one left-to-right and one right-to-left phase encoding direction acquired on each day.

### UK Biobank

To investigate the association between prediction accuracy and reliability as well as how reliability interacts with sample size, we randomly sampled n = 5000 (2714 female, ages = 48-82) participants from healthy subjects of the UK Biobank sample (Sudlow et al., 2015). Healthy participants were defined as subjects without lifetime prevalence of cerebrovascular diseases, infectious diseases affecting the nervous system, neuropsychiatric disorders or neurological diseases based on ICD-10 diagnosis from hospital inpatient records and self-report (see supplementary methods for all excluded data fields). All subjects had complete rs-fMRI scans and displayed no excessive head movement (framewise displacement < 0.28 mm, which corresponded to a displacement of 3SD above the mean). Within this sample, we selected 17 phenotypes that were available for all subjects and did not display strong ceiling effects (see supplementary methods for a full list of phenotypes and their distributions). Of those, age and grip strength were used for creating simulated data. Additionally, a sample of 1890 (1012 female, ages = 48-79) subjects with available follow-up data for all 17 phenotypes from the follow-up imaging session was used to estimate phenotypic reliability. The mean interval between the initial imaging session and the follow-up session was 2 years and 6 months.

The UKB scanning protocol (Miller et al., 2016) included structural and resting state fMRI images acquired at four imaging centres (Bristol, Cheadle Manchester, Newcastle and Reading) with harmonized Siemens 3T Skyra MRI scanners with a 32-channel head coil. T1w MRI images were acquired using a 3D MPRAGE sequence (TR = 2000 ms, 1.0 mm isotropic). One rs-fMRI session with 490 volumes each (6 min and 10 s) was acquired using a multiband echo-planar imaging (TR = 735 ms, 2.4 mm isotropic voxels).

### Adolescent Brain Cognitive Development

To investigate if our association between phenotype reliability and prediction accuracy generalises to an additional dataset with different preprocessing we used data from the Adolescent Brain Cognitive Development study (Volkow et al., 2018) baseline sample from the ABCD BIDS Community Collection (Feczko et al., 2021). Only English speaking participants without severe sensory, intellectual, medical or neurological issues and all available behavioural phenotypes were used (see supplementary methods for a full list). Furthermore, all participants had to have complete rs-fMRI data and pass the ABCD quality control for their T1 and resting-state fMRI. This resulted in a total of 4133 participants (2123 female, ages = 9-11). Additionally, a sample of 2102 (1026 female, ages = 9-11) subjects with available follow-up data for all phenotypes from the first follow-up session was used to estimate phenotypic reliability. The mean interval between the initial imaging session and the follow-up session was 1 year and 11 months.

The ABCD acquisition protocol (Casey et al., 2018) was harmonised across 21 sites on Siemens Prisma, Phillips, and GE 750 3T scanners. It included high-resolution T1w MRI images with a 32-channel head coil using a 3D multi-echo MPRAGE sequence (TR = 2500 ms, 1.0 mm isotropic voxels). The rs-fMRI images were acquired using gradient-echo echo-planar imaging (TR = 800 ms, 2.4 mm isotropic voxels) and included two sessions totalling 20 minutes.

### Simulation of different levels of reliability of selected phenotypes

For each of the selected prediction targets we created simulated datasets with varying amounts of noise. According to classical measurement theory (Novick, 1966), any measurement reflects a mixture of the measured entity and random (as well as systematic) measurement noise. The reliability of a variable can thus be reduced by increasing the proportion of error or noise variance while holding between-subject variance constant, thereby reducing the signal-to-noise ratio. Here we manipulated only the unsystematic measurement noise, defined as random variability that produces a discrepancy between observed and true values (or repeated observations). Increasing random noise is ideal for investigating test-retest reliability as it only affects the variability of measurements around the mean and thus manipulates the ranking across individuals.

In order to induce increasing levels of noise in the target variable, we created datasets that correlated with the originally observed (empirical) targets at a pre-specified Pearson’s correlation. This method was chosen to increase the interpretability of the resulting attenuation of brain-behaviour associations by controlling the amount of noise. The data generation procedure was as follows: First, a random vector was sampled from a standard normal distribution with the same mean and standard deviation as the original empirically acquired data (in the HCP these were age-adjusted and normalised to mean = 100 and SD = 15). Next, we calculated the residuals of a least squares regression of the sampled vector (X) on the empirical data (Y). The resulting orthogonal vector representing the portion of X that is independent of Y was then again combined with the original empirical data Y through scaling by the pre-specified correlation. This adjustment process manipulated the relative contributions of Y and the residuals of X on Y in the resulting simulated vector. The formula used for this process was:

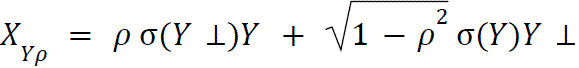

Where X_Y*Q*_ is the new ‘simulated’ vector that correlates with the empirical data Y at a predefined correlation ρ. *Y* ⊥ represents the residuals of a least squares regression of a randomly sampled vector X against Y. We provide an R implementation in the accompanying repository online (https://github.com/MartinGell/Reliability/code).

The pre-specified correlations for simulated data based on the HCP-A dataset were set to correlate with the original data at r = 0.99, 0.95, 0.9, 0.85, 0.8, 0.75, 0.7, 0.65, 0.6, 0.55 and 0.5. Given the high computational load for large samples, simulated UKB data were set to correlate at r = 0.9, 0.8, 0.7, 0.6 and 0.5 with the original data. For each level of correlation, simulation was repeated 100 times, thus totalling 4400 simulated datasets for HCP-A (4 assessed phenotypes x 11 noise levels x 100 repeats) and 1500 simulated datasets for UKB (3 assessed phenotypes x 5 noise levels x 100 repeats). Simulated datasets were scaled and offset to have approximately the same mean and standard deviation as the original measurements to facilitate absolute agreement (i.e. stability across repeated measurements) between original data and the simulated test-retest data in order to harmonise test-retest correlations and ICC (for distribution of mean and SD for each dataset see the accompanying online repository: https://github.com/MartinGell/Reliability/plots). As age did not follow a normal distribution, we first estimated its probability density from the original data and then sampled simulated data from this distribution instead.

### Phenotype preprocessing

As we used linear ridge regression for prediction, all phenotypes that displayed a right-skewed distribution were transformed with a natural log transform. As this procedure manipulated data within participants, there was no data leakage across participants.

### fMRI Preprocessing

Both HCP datasets provided minimally preprocessed data. The preprocessing pipeline has been described in detail elsewhere (Glasser et al., 2013). Briefly, this included gradient distortion correction, image distortion correction, registration to subjects’ T1w image and to MNI standard space followed by intensity normalisation of the acquired rs-fMRI images, and Independent Component Analysis (ICA) followed by an ICA-based X-noiseifier (ICA-FIX) denoising (Beckmann & Smith, 2004; Salimi-Khorshidi et al., 2014) Additional denoising steps were conducted by regressing mean time courses of white matter and cerebrospinal fluid and the global signal, which has been shown to reduce motion-related artefacts (Ciric et al., 2017). Next, data were linearly detrended and bandpass filtered at 0.01 – 0.1 Hz.

The UKB data were preprocessed through a pipeline developed and run on behalf of UK Biobank (Alfaro-Almagro et al., 2018) and included the following steps: motion correction using MCFLIRT (Jenkinson, Bannister, Brady, & Smith, 2002); grand-mean intensity normalisation of the entire 4D fMRI dataset by a single multiplicative factor; highpass temporal filtering using Gaussian-weighted least-squares straight line fitting with sigma = 50 sec; Echo Planar Imaging unwarping; Gradient Distortion Correction unwarping; structured artefact removal through ICA-FIX (Beckmann & Smith, 2004; Salimi-Khorshidi et al., 2014). No low-pass temporal or spatial smoothing was applied. The preprocessed datasets (named as *filtered_func_data_clean.nii* in the UK Biobank database) were normalised to MNI space using FSL’s *applywarp* command.

The ABCD dataset was preprocessed ABCD-BIDS pipeline as part of the ABCD-BIDS Community Collection (ABCC; Collection 3165) which has been described in detail elsewhere (Feczko et al., 2021). The pipeline included distortion correction and alignment using Advanced Normalization Tools (ANTS), FreeSurfer segmentation, and surface as well as volume registration using FSL FLIRT rigid-body transformation. Processing was done according to the DCAN BOLD Processing (DBP) pipeline which included de-trending and de-meaning of the rs-fMRI data, denoising using a general linear model with regressors for tissue classes and movement. The data were then bandpass filtered between 0.008 and 0.09 Hz using a 2nd order Butterworth filter. DPB respiratory motion filtering (18.582 to 25.726 breaths per minute), and censoring (frames exceeding an FD threshold of 0.2mm or failing to pass outlier detection at +/− 3 standard deviations were discarded) were then applied.

### Functional connectivity

The denoised time courses from all datasets were parcellated using the Schaefer et al. (2018) atlas with 400 cortical regions of interest for all main analyses. The signal time courses were averaged across all voxels of each parcel. Parcel-wise time series were used for calculating functional connectivity between all parcels using Pearson correlation. For HCP datasets, the correlation coefficients of individual sessions (4 per participant) were transformed into Fisher-Z scores, and for each connection, an average across sessions was calculated. To investigate the robustness of our results to granularity and parcellation selection, functional connectivity between denoised time courses of 200, 300 cortical regions from the Schaefer et al. (2018) atlas as well as 300 cortical, subcortical and cerebellar regions of interest defined by Seitzman et al. (2020) was calculated. Regions were modelled as 6-mm spheres and calculated from resting state data from the HCP Aging dataset (results presented in the supplemental material). Finally, to investigate the generalisation of our results to another dataset with different preprocessing steps, the ABCD dataset was parcellated using HCP’s 360 ROI atlas template (Glasser et al., 2016).

### Prediction

We used the scikit-learn library [version 0.24.2, (Pedregosa et al., 2011)] to predict all target variables from functional connectivity (code including exemplary data available online: https://github.com/MartinGell/Prediction_Reliability). Accuracy was measured using R^2^ (coefficient of determination), mean absolute error (MAE) and Pearson correlation between predicted and observed target values. The R^2^ represents the proportion of variance (in the target variable) that has been explained by the independent variables in the model and is calculated here as

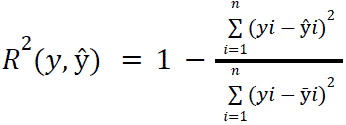

Where 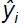 is the predicted value of the i-th sample and *y_i_* is the corresponding true value for total n samples. 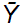 Represents the mean across all *y*. In this formulation, the R^2^ is not interchangeable with the correlation coefficient squared. All predictions were performed using linear ridge regression as it showed a favourable ratio of computation time to accuracy in previous work (Cui & Gong, 2018) and preliminary testing (see supplementary methods).

Out-of-sample prediction accuracy was evaluated using a nested cross-validation with 10 outer folds and 5 repeats. Hyperparameter optimization (inner training folds) of the α regularisation parameter for ridge regression was done using efficient leave-one-out cross-validation (Rifkin & Lippert, 2007). The model with the best α parameter was then fitted on the training folds and tested on the outer test folds. Within each training fold, neuroimaging features were standardised by z-scoring across participants before models were trained in order to ensure that individual features with large variance would not dominate the objective function. Before prediction (of both original and simulated data), subjects with target values 3 SD from the sample mean were removed from the complete sample to minimise the impact of extreme values resulting from random sampling in simulated data. As a preprocessing step prior to training, neuroimaging features were z-scored within participants.

### Control analyses for simulation results in HCP-A

To verify our analyses were robust to analytical degrees of freedom, we repeated our analyses of the HCP-A dataset using support vector regression, an alternative node definition for functional connectivity features (using ROIs from Seitzman et al., 2020) and feature-wise confound removal. For algorithm comparison, we trained a support vector regression with a linear kernel on neuroimaging features. Out-of-sample prediction accuracy was evaluated using a non-nested cross-validation with 10 outer folds and 5 repeats. A heuristic was used to efficiently calculate the hyperparameter C (Helleputte, Paul, & Gramme, 2021):

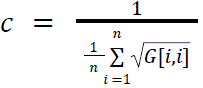

where G is the matrix multiplication of features and transposition of features (here: functional connectivity). To investigate whether confounding effects impacted our results, standard confound variables (age and sex for the prediction of all phenotypes) were removed from the connectivity features using linear regression. Confound removal was performed within each training fold and the confound models were subsequently applied to test data to prevent data leakage (More, Eickhoff, Caspers, & Patil, 2021). The impact or the reliability of functional connectivity was investigated by reducing the length of the resting state time courses before calculating functional connectivity. Firstly, only two sessions (rather than all four), corresponding to the resting-state images acquired on the first day, were used. Finally to mirror the UKB acquisition protocol only the very first session acquired in the anterior-to-posterior direction was used for calculating functional connectivity. All control analyses are presented in the supplemental material.

### Association between reliability and prediction accuracy

The relationship between target reliability and prediction accuracy (measured as R^2^) was investigated using the HCP-YA dataset. First, the test-retest data of 46 participants was used to estimate measurement reliability for 36 different behavioural phenotypes by calculating ICC between the scores from the first and second visits. ICC was calculated using a two-way random effects model for absolute agreement, often referred to as ICC[2,1] (Shrout & Fleiss, 1979). Next, all selected measures were predicted in a sample of 713 subjects from the HCP-YA dataset using linear ridge regression. As the HCP-YA dataset includes related subjects, cross-validation was done using a 5 times repeated leave 30% of families out approach, instead of the 10-fold random split used in other analyses. Family members were always kept within the same fold in order to maintain independence between the folds. Confounding effects of age and sex on features were removed using linear regression trained on the training set and applied to test data within the cross-validation. Finally, the resulting prediction accuracies (R^2^) of the 36 different phenotypes were correlated with their corresponding reliability (calculated from the test-retest data). To validate our findings, the above-described approach (with the exception of cross-validation) was repeated using the UKB dataset. Reliability was estimated for 17 different behavioural assessments using ICC2 between measurements collected during the first and follow-up imaging visits in 1893 subjects. All phenotypes were predicted in a set of 5000 subjects from the UKB using ridge regression in nested cross-validation with 10 outer folds and 5 repeats used for our main analyses.

### Subsampling procedure and prediction in the UKB dataset

To examine how the effects of reliability on prediction performance interact with increasing sample size, we randomly sampled geometrically spaced samples (series with a constant ratio between successive elements) from 5000 subjects of the UK Biobank starting from n = 250 (250, 403, 652, 1054, 1704, 2753, 4450). By doing so, we aimed to cover sample sizes ranging from those available in larger neuroimaging studies to international consortia levels. To be able to compare prediction accuracy between different sample sizes we used a learning curve function from Sklearn (‘learning_curv’). In this approach, we first partitioned a test set of 10% of the full sample (500 subjects). From the remaining data, geometrically spaced samples of subjects (250, 403, 652, 1054, 1704, 2753, 4450) were sampled without replacement. Each subsample was then used to train a ridge regression model with hyperparameter optimization using the same cross-validation set-up with 10 outer folds and 5 repeats used in previous analyses. This approach made the comparison of accuracy between different sample sizes possible as the test set is held constant for all samples of training subjects. The entire procedure was repeated 100 times for all simulated and empirical data.

## Supporting information

Supplementary Results Figure 1-16

## Acknowledgements

This research has been conducted using the UK Biobank Resource under Application Number 41655. Funding was provided by the Deutsche Forschungsgemeinschaft (DFG, German Research Foundation) – 269953372/GRK2150

