## Supplementary Results Figure 1-16 for "The Burden of Reliability: How Measurement Noise Limits Brain-Behaviour Predictions"

#### Prediction accuracy for age prediction - additional metrics

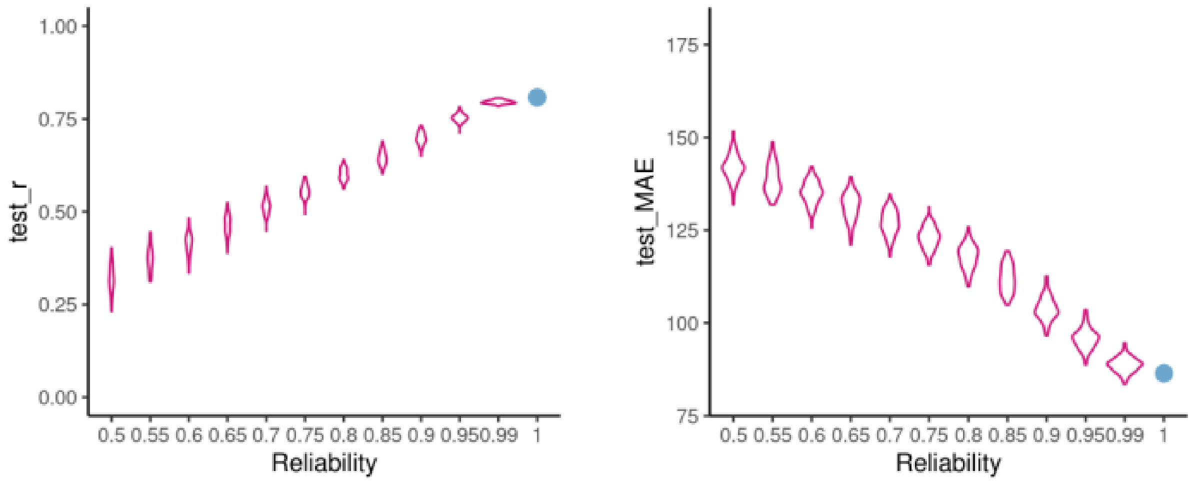


**Supplementary Figure 1.** Mean absolute error (MAE) and correlation between observed and predicted targets in age prediction

#### Sensitivity analyses - Age prediction

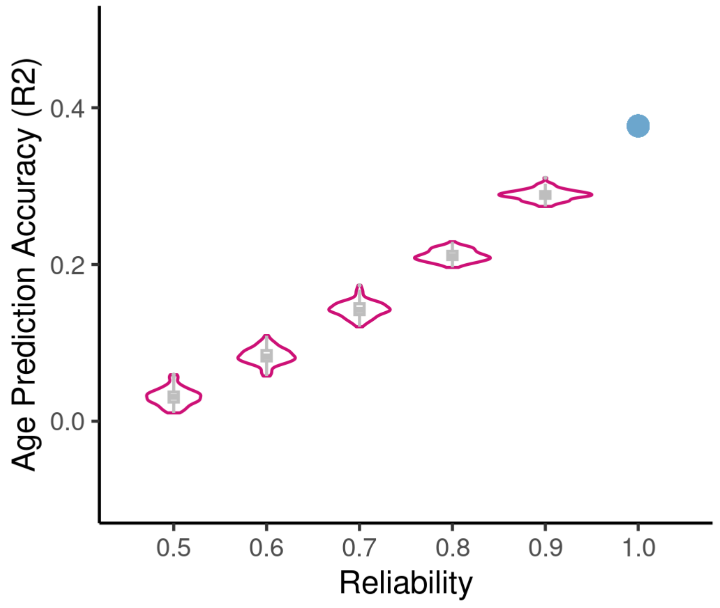


**Supplementary Figure 2.** Prediction of age using 5000 subjects from the UKB. The same sample of participants was used to create this figure as in the section “Behavioural reliability

is related to prediction accuracy” of the results. The age of subjects was predicted using ridge regression as in Figure 1 and accuracy was evaluated using R2.

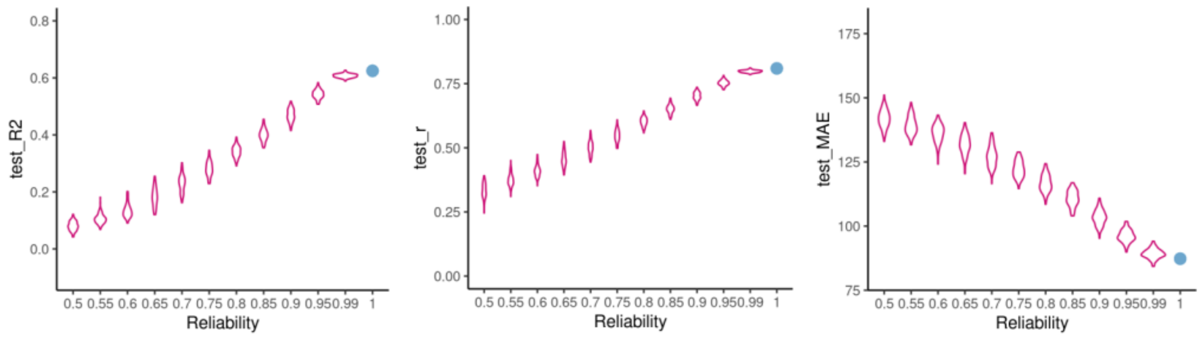


**Supplementary Figure 3.** Prediction Using Seitzman et al. (2020) Nodes. R2, Mean absolute error (MAE) and correlation between observed and predicted targets in age prediction using 300 nodes by Seitzmann et al. 2020

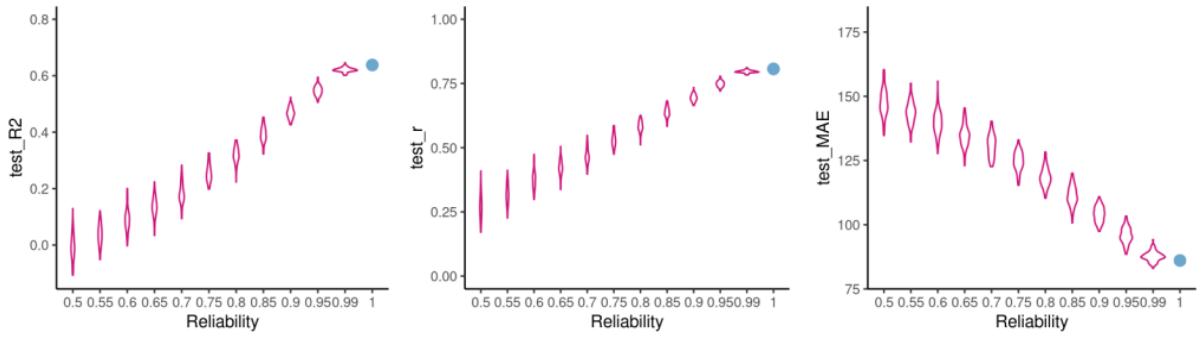


**Supplementary Figure 4.** Prediction Using Support Vector Regression. R2, Mean absolute error (MAE) and correlation between observed and predicted targets in replication of age prediction using support vector regression

#### Impact of reliability on prediction accuracy in the HCP-A dataset - influence of connectivity reliability.

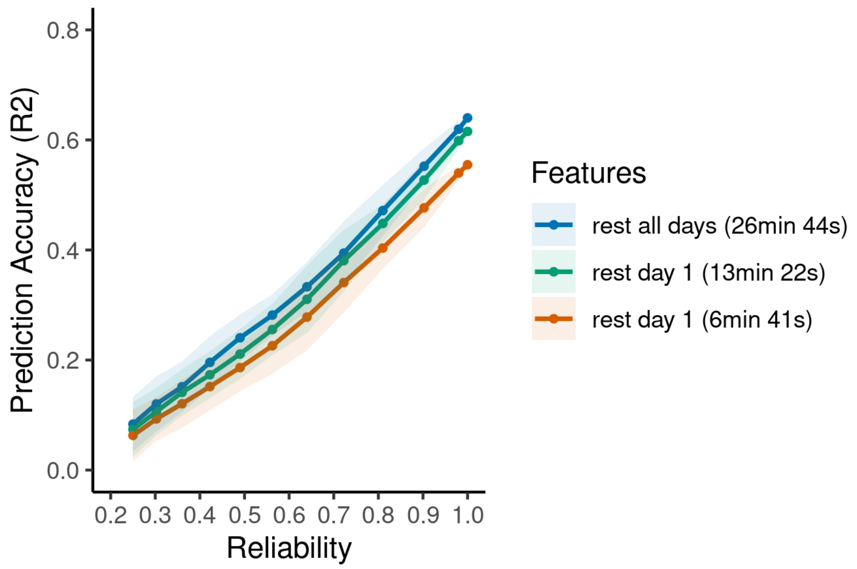


**Supplementary Figure 5.** Influence of feature reliability (functional connectivity) on age prediction with decreasing reliability. Impact of shortening rsfMRI timecourses from the average of all four sessions collected on both days to an average of 13 minutes collected on the first day only and finally connectivity calculated from a single 6 minutes session collected on the first day in the anterior-to-posterior direction (same as only session in the UKB).

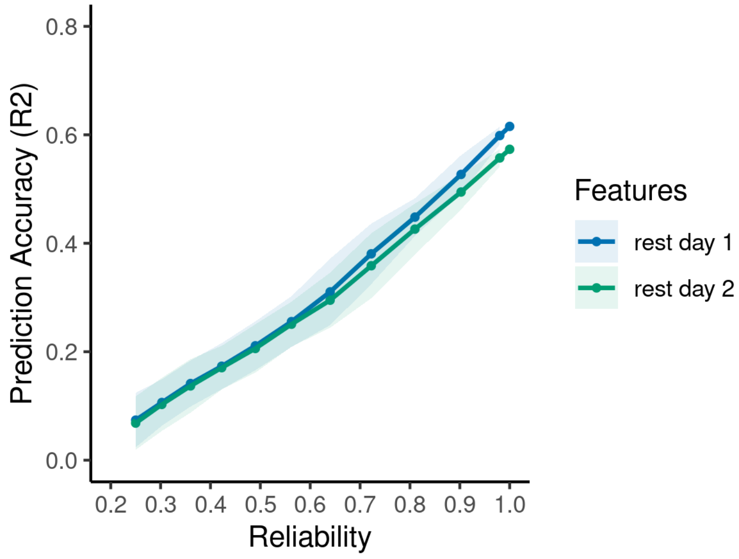


**Supplementary Figure 6.** Influence of feature reliability (functional connectivity) on age prediction with decreasing reliability. Age prediction from day 1 and 2 rsfMRI acquisition separately. Both days had 13 minutes and 22 seconds of data.

#### Sensitivity analyses - prediction of behaviour

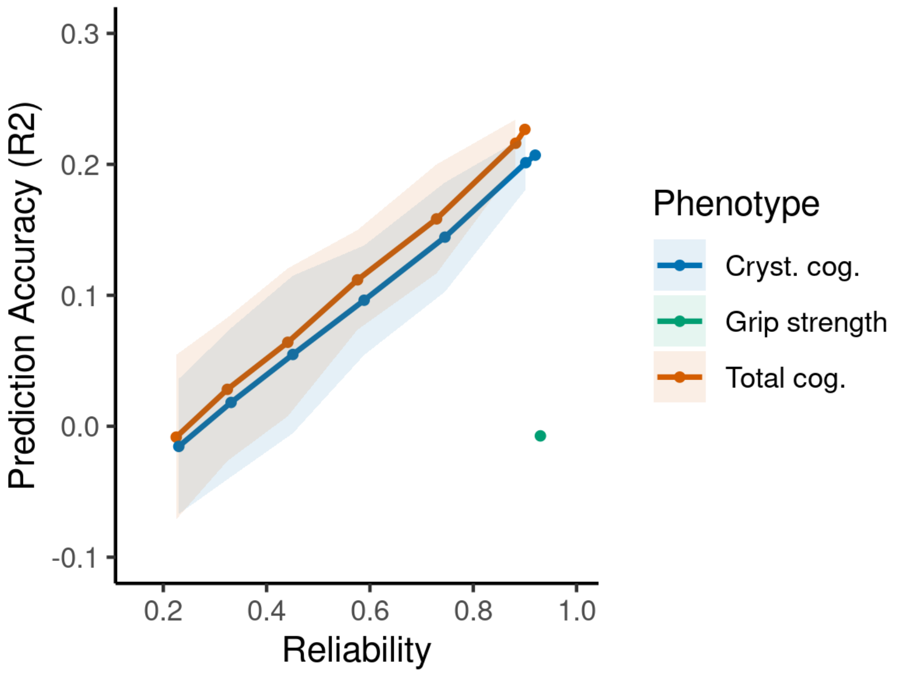


**Supplementary Figure 7.** Prediction of selected phenotypes with feature-wise confound (age, sex) regression. Grip strength could not be predicted when confounds were regressed.

#### Sensitivity analyses - Total cognition

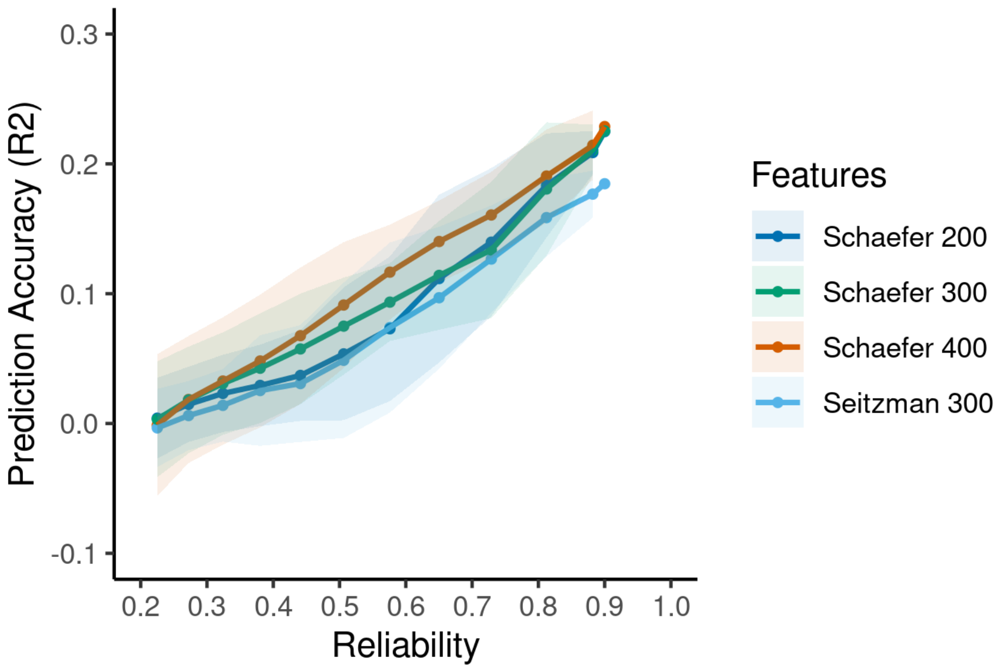


**Supplementary Figure 8.** Prediction of selected phenotypes with feature-wise confound (age, sex) regression. Grip strength could not be predicted when confounds were regressed.

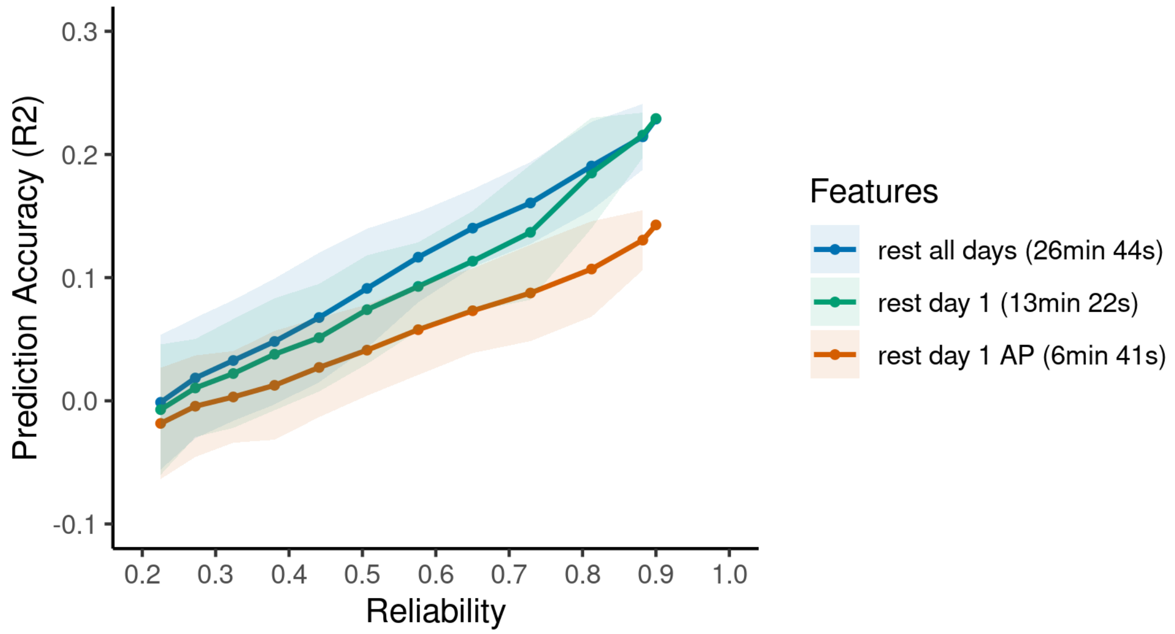


**Supplementary Figure 9.** Influence of feature reliability (functional connectivity) on prediction of total cognition with decreasing reliability. Impact of shortening rsfMRI timecourses from the average of all four sessions collected on both days to an average of 13 minutes collected on the first day only and finally connectivity calculated from a single 6 minutes session collected on the first day in the anterior-to-posterior direction (same as only session in the UKB).

### Phenotype prediction supplementary figures

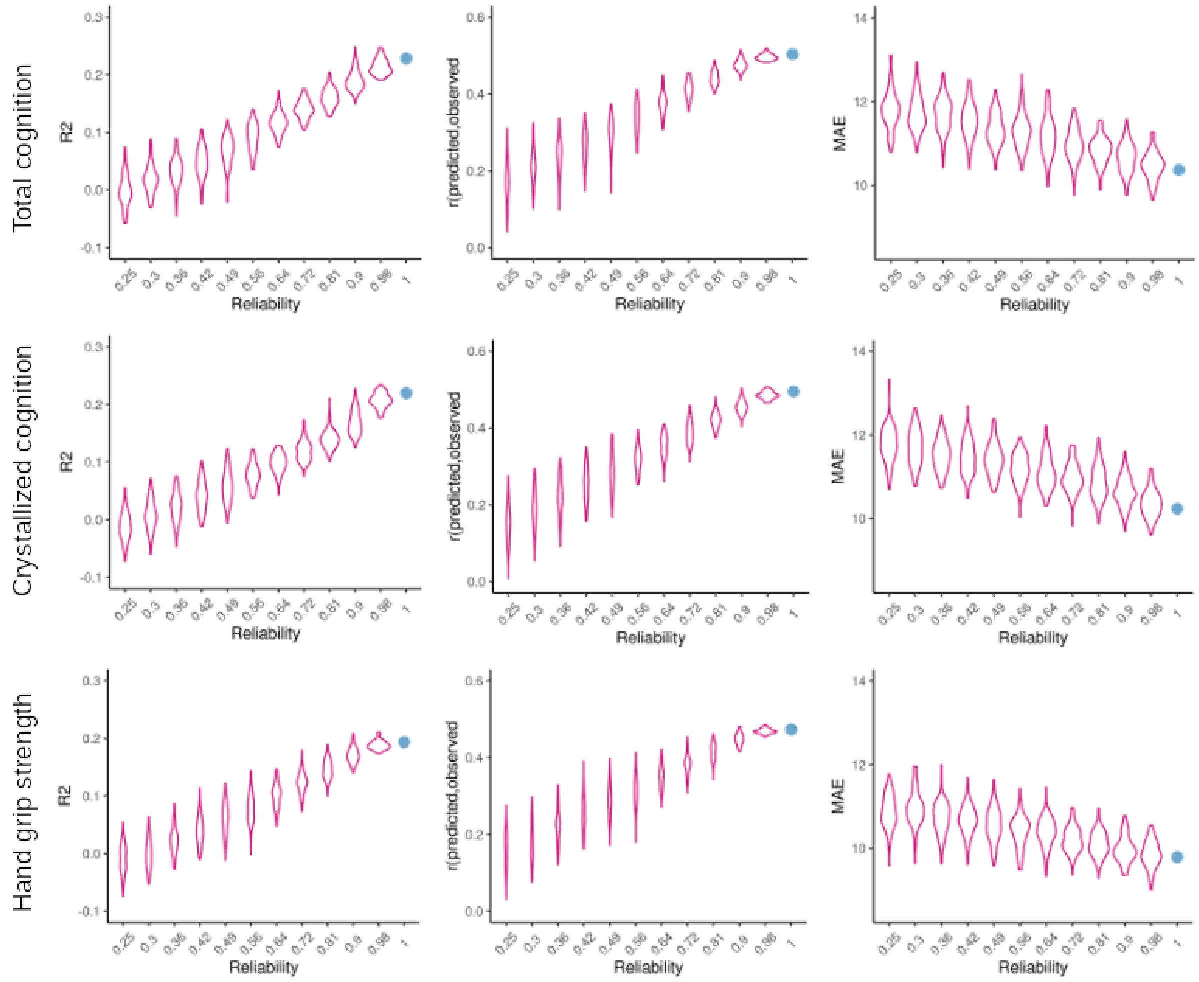


**Supplementary Figure 10.** R2, Mean absolute error (MAE) and correlation between observed and predicted targets for each target behaviours separately.

#### Uncorrected results for Figure 1

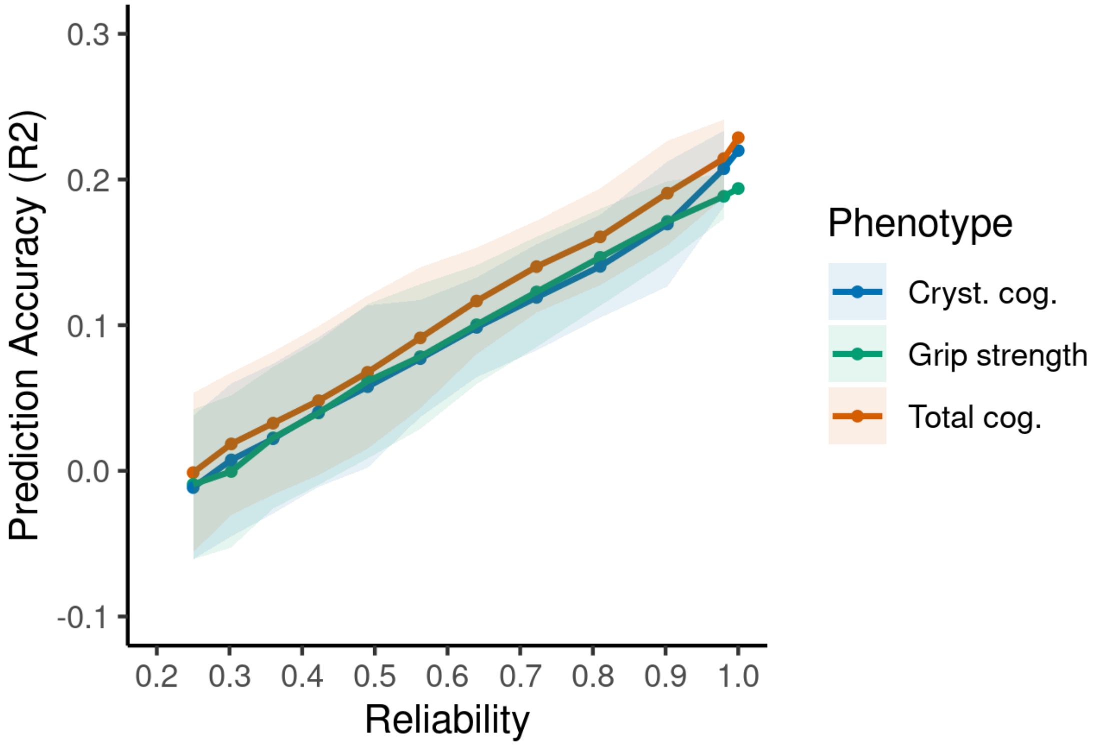


**Supplementary Figure 11.** Impact of reducing the correlation between original and simulated target scores (reflecting reduced reliability) on accuracy in prediction of total cognition composite score, crystallised cognition composite score and grip strength. Solid lines represent the mean across all 100 simulated datasets in each correlation band, shaded areas represent 2 standard deviations in prediction accuracies.

#### HCP young adult dataset predicted behaviours and respective reliability

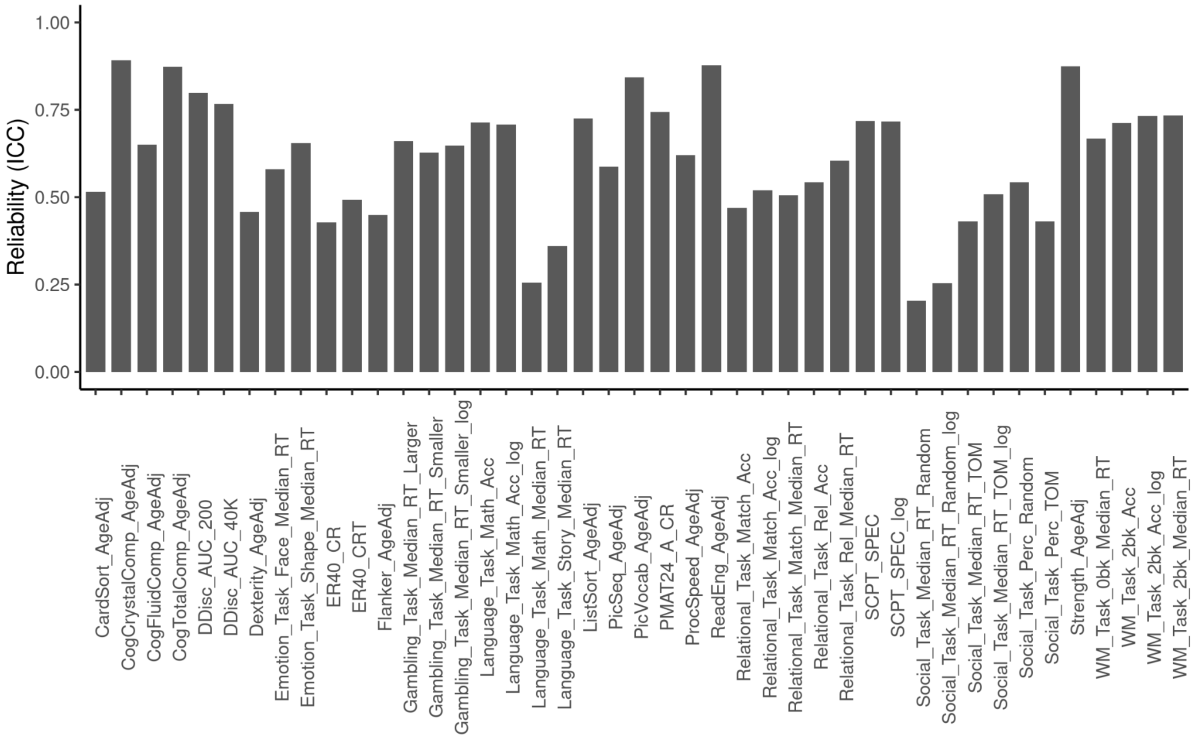


**Supplementary Figure 12.** Reliability of all behaviours predicted in the HCP-YA dataset

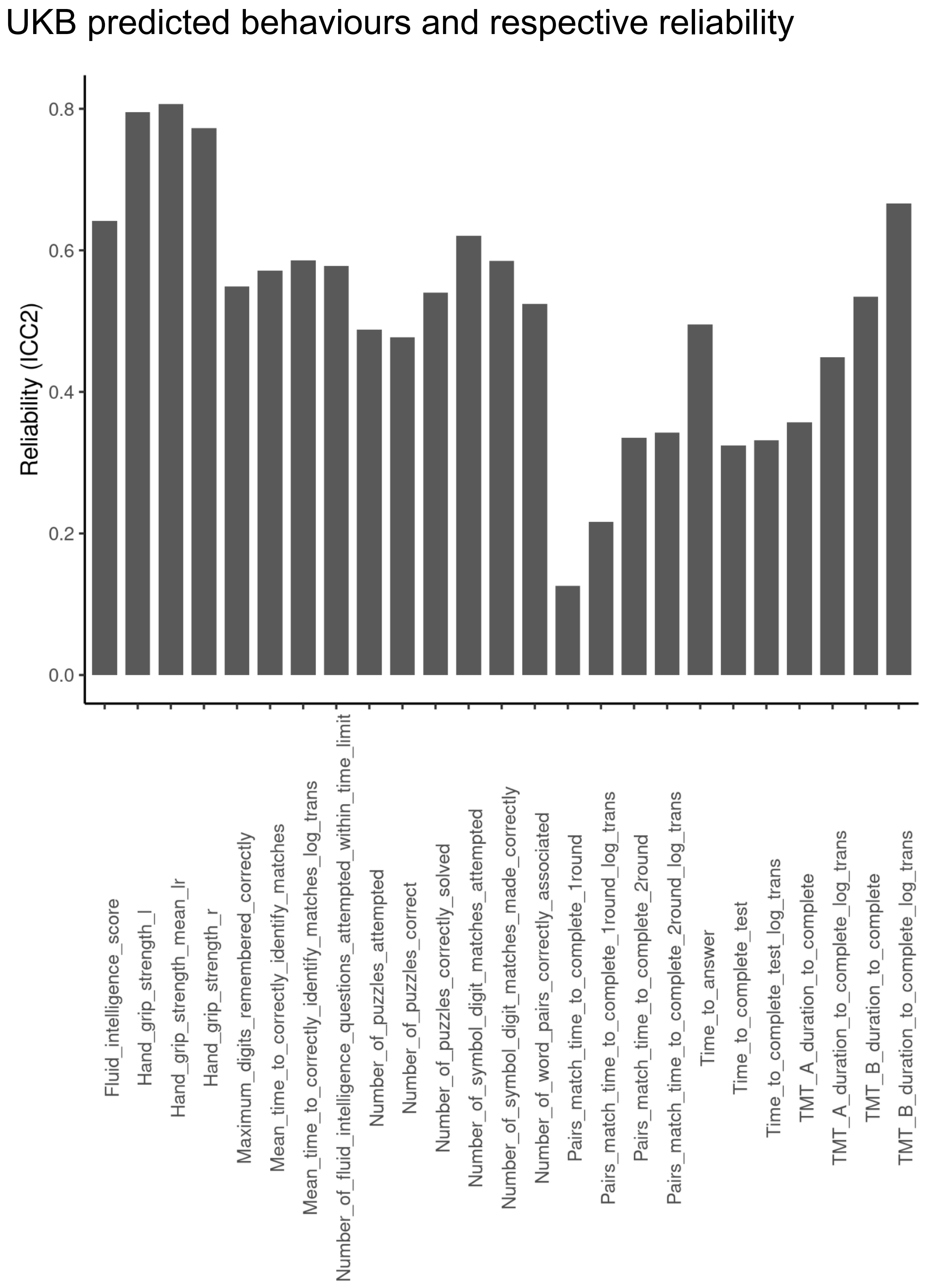


**Supplementary Figure 13.** Reliability of all behaviours predicted in the UKB dataset

#### Association between prediction accuracy and reliability in the ABCD dataset

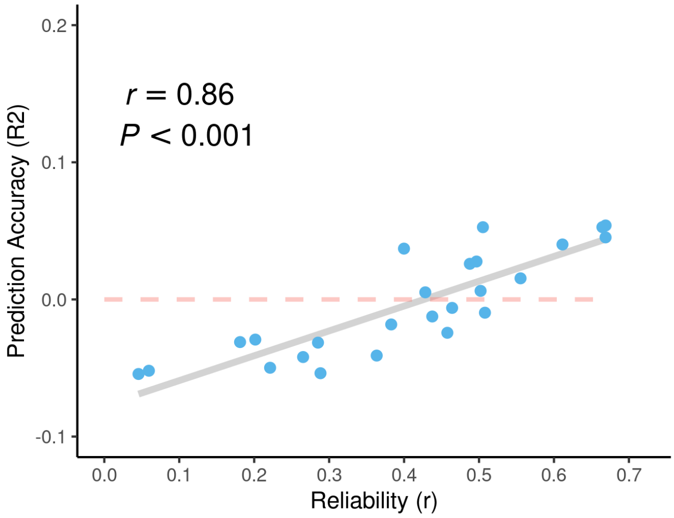


**Supplementary Figure 14.** Association between reliability and prediction accuracy. Each data point represents one of 25 behavioural assessments in the ABCD. As measurements were collected from participants aged 9-10 at baseline and 11-12 at follow-up, reliability is calculated with test-retest (Pearson) correlation and not ICC. Unlike ICC, correlation is robust to systematic age-related changes as it is not penalised by differences in means between baseline and follow-up data (mean retest interval = 23 months) and different rates of development across participants.

#### Prediction of cognitive flexibility measured trail-making task in UKB

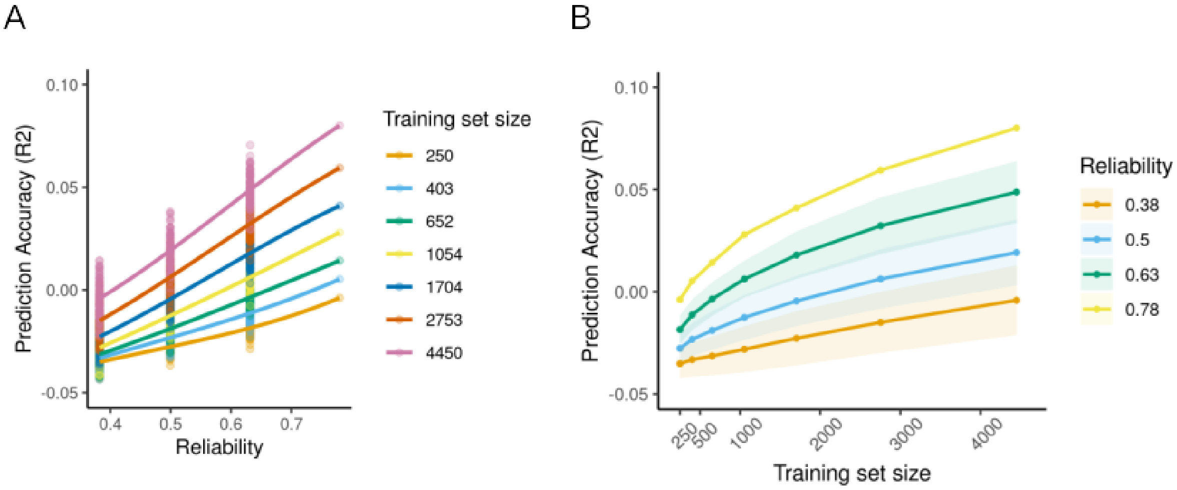


**Supplementary Figure 15.** Prediction and subsampling in UKB. (A) Impact of training set size on original and simulated cognitive flexibility with reduced reliability. Panel (B) Impact of training set size on prediction accuracy in empirical and simulated data with varying levels of reliability. Solid lines represent the mean across all 100 simulated datasets in each correlation band and shaded areas represent 2 standard deviations in prediction accuracies. Results were fitted with an exponential function for illustration purposes and were adjusted for the reliability of the TMT task (ICC = 0.775) estimated in independent data by Fawns-Ritchie et al. (2020).

#### Learning curve results for empirical behaviours

**Supplementary Results Table 1**

*Prediction accuracy (R2) of empirical behaviours across varying training set sizes*

| Behaviour | Training Set Size | | | | | | |
|---|---|---|---|---|---|---|---|
| | 250 | 403 | 652 | 1054 | 1704 | 2753 | 4450 |
| Fluid Intelligence | -0.02 | -0.014 | -0.006 | 0.004 | 0.008 | 0.018 | 0.029 |
| Associative Learning (SDST) | -0.001 | 0.009 | 0.02 | 0.033 | 0.043 | 0.056 | 0.068 |
| Cognitive Flexibility (TMT B) | -0.002 | 0.007 | 0.0198 | 0.03 | 0.042 | 0.056 | 0.069 |
| Hand Grip Strength (mean) | 0.127 | 0.16 | 0.197 | 0.24 | 0.279 | 0.313 | 0.344 |
| Age | 0.107 | 0.149 | 0.189 | 0.228 | 0.272 | 0.316 | 0.354 |

#### Improvement in prediction accuracy with increased training set size

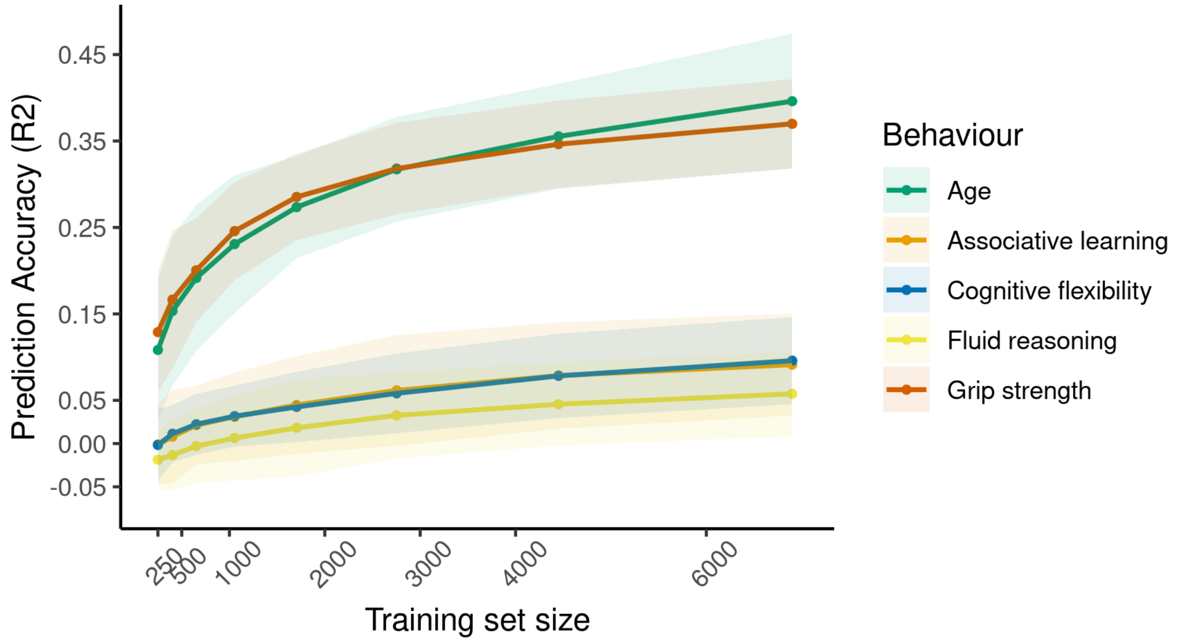


**Supplementary Figure 16.** Impact of training set size on prediction accuracy of empirical behaviours. Solid lines represent the mean accuracy across 100 subsamples and shaded areas represent 2 standard deviations in prediction accuracy. Abbreviations: SDST, Symbol Digit SubstitutionTest; TMT-B, Trail making task part B.

Distribution of test-retest correlations across HCP-YA, UK and ABCD datasets

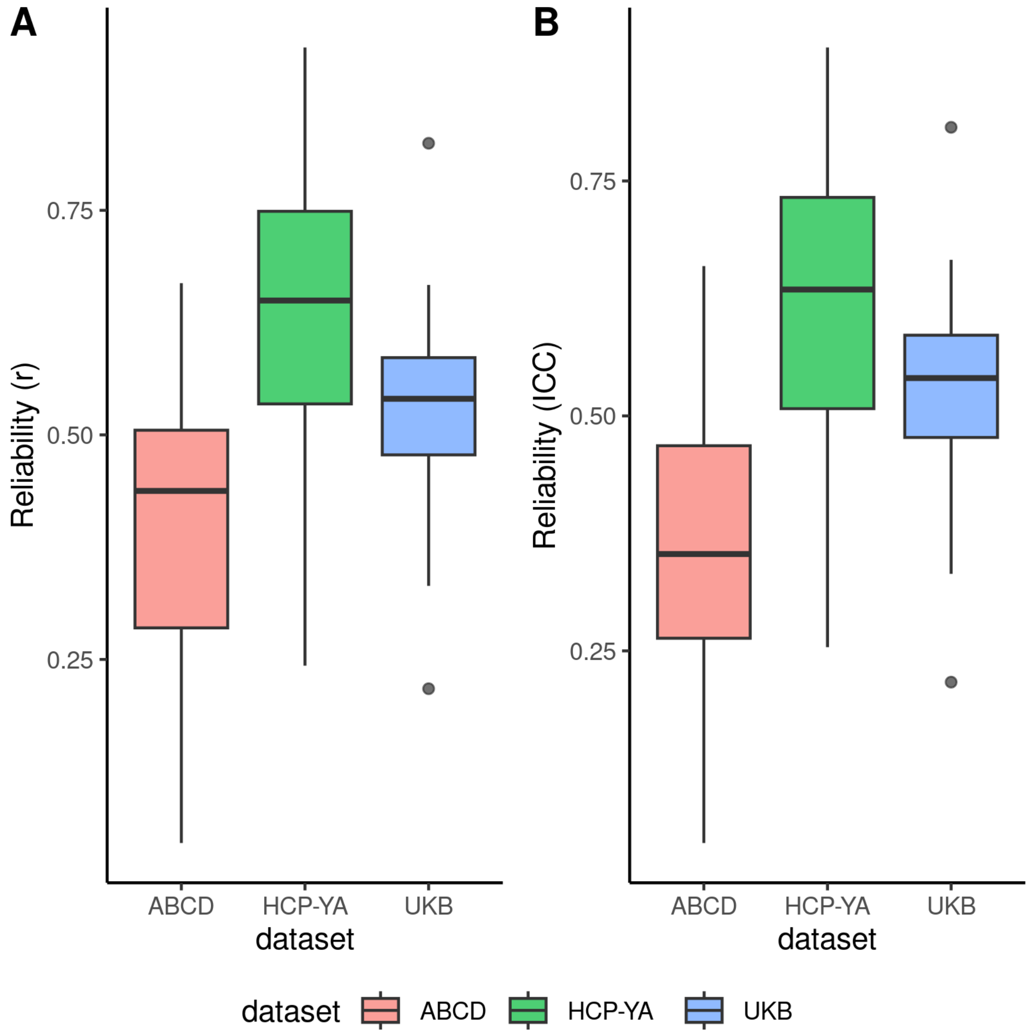


**Supplementary Figure 17.** Boxplots of test-retest correlations (A) and ICC (B) of all measures used for assessing the association between prediction accuracy and reliability in Figure 2 of the main text. Overall median for test-retest correlations = 0.55; and ICC = 0.51. Both retest correlations and ICC are displayed as for some datasets, such as the ABCD, test-retest correlations may be more appropriate as systematic error coming from different rates of development across participants is not penalised.

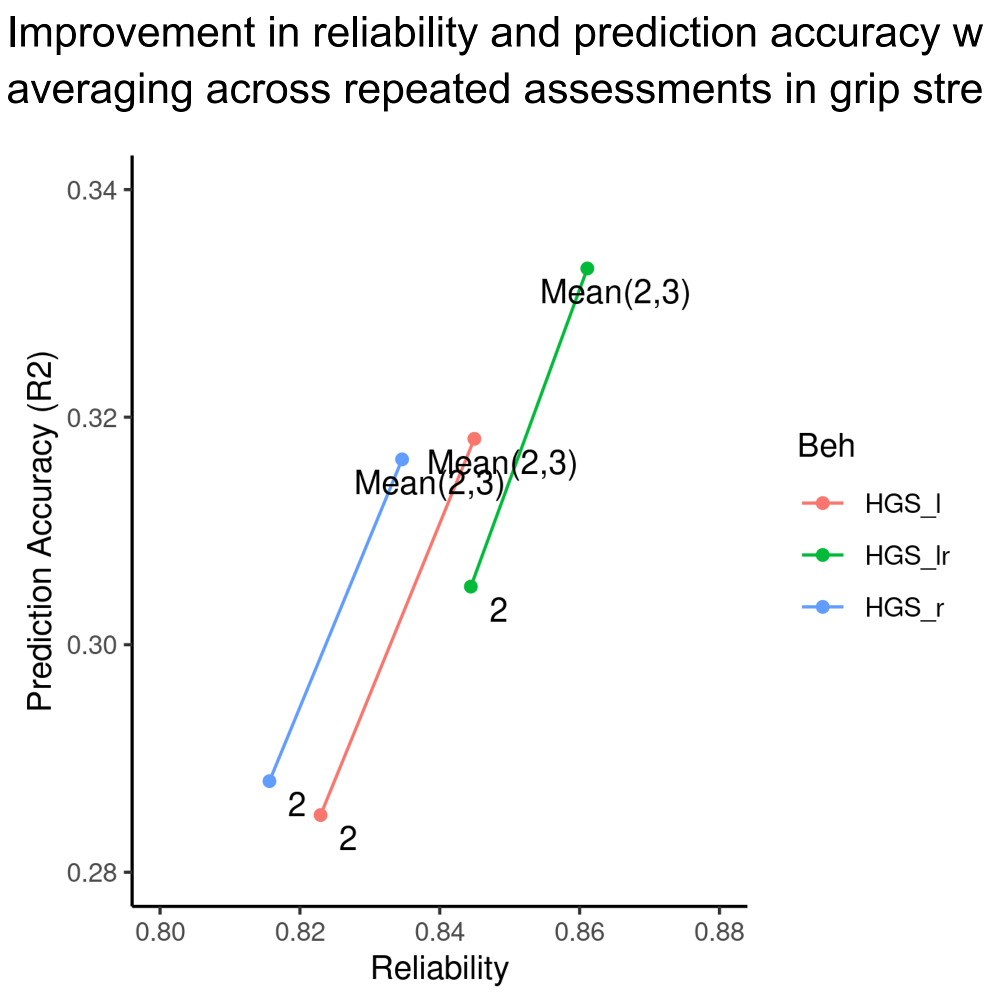


**Supplementary Figure 18.** Impact of increasing reliability by averaging in UKB hand grip strength prediction. Numbers denote UKB time points. Neuroimaging was collected at time point 2. Grip strength was averaged over time points 2 (neuroimaging baseline) and 3 (neuroimaging follow-up) to maximise the number of subjects. Reliability was calculated as Pearson test-retest correlation between time point 0 (baseline) and time point 2 as well as the average of 2 and 3. HGS_lr stands for the average of measurement over left and right hands. Abbreviations; HGS: Hand grip strength.
