## Supplementary Methods for "The Burden of Reliability: How Measurement Noise Limits Brain-Behaviour Predictions"

Table 1 and 2 display all predicted behaviours and accompanying figures 1 and 2 display the data distributions in HCP-YA and UKB respectively. The next section details excluded and included fields in parsing UKB subjects. Table 3 contains an overview of exact sample size for HCP-A subjects. The last section details our algorithm comparison.

#### 1. Behaviours selected for prediction in HCP-YA dataset

**Supplementary Table 1**

*Predicted behaviours in HCP-YA*

| Behaviour |
|---|
| PicSeq_AgeAdj |
| CardSort_AgeAdj |
| Flanker_AgeAdj |
| PMAT24_A_CR |
| ReadEng_AgeAdj |
| PicVocab_AgeAdj |
| ProcSpeed_AgeAdj |
| DDisc_AUC_200 |
| DDisc_AUC_40K |
| ListSort_AgeAdj |
| SCPT_SPEC |
| CogFluidComp_AgeAdj |
| CogTotalComp_AgeAdj |
| CogCrystalComp_AgeAdj |
| ER40_CR |
| ER40_CRT |

Strength_AgeAdj

Dexterity_AgeAdj

WM_Task_2bk_Acc

WM_Task_2bk_Median_RT

WM_Task_0bk_Median_RT

Language_Task_Math_Acc

Language_Task_Math_Median_RT

Language_Task_Story_Median_RT

Social_Task_Perc_Random

Social_Task_Perc_TOM

Social_Task_Median_RT_Random

Social_Task_Median_RT_TOM

Relational_Task_Match_Acc

Relational_Task_Match_Median_RT

Relational_Task_Rel_Acc

Relational_Task_Rel_Median_RT

Emotion_Task_Face_Median_RT

Emotion_Task_Shape_Median_RT

Gambling_Task_Median_RT_Larger

Gambling_Task_Median_RT_Smaller

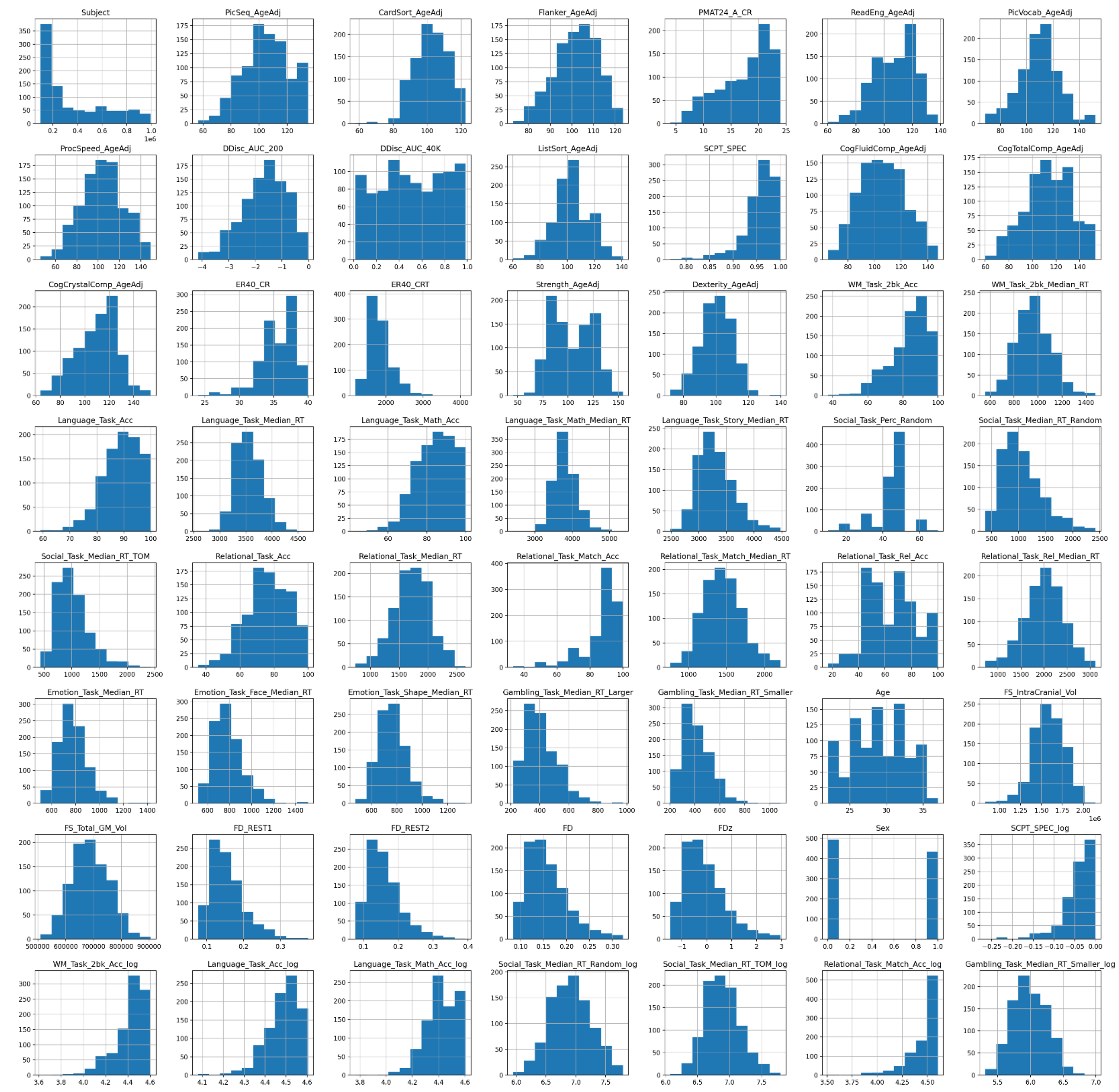


**Supplementary Figure 1.** Distribution of all predicted behaviours HCP-YA

### 2. Behaviours selected for prediction in UKB sample

**Supplementary Table 2**

*Predicted behaviours in UKB*

| Behaviour | Data-field |
|---|---|
| Pairs_match_time_to_complete_1st_round-2.0 | 400 |
| Pairs_match_time_to_complete_2nd_round-2.0 | 400 |
| Maximum_digits_remembered_correctly-2.0 | 4282 |
| Time_to_complete_test-2.0 | 4285 |
| Time_to_answer-2.0 | 4288 |
| TMT_A_duration_to_complete-2.0 | 6348 |
| TMT_B_duration_to_complete-2.0 | 6350 |
| Number_of_puzzles_correctly_solved-2.0 | 21004 |
| Number_of_puzzles_attempted-2.0 | 6383 |
| Fluid_intelligence_score-2.0 | 20016 |
| Mean_time_to_correctly_identify_matches-2.0 | 20023 |
| Number_of_fluid_intelligence_questions_attempted_within_time_limit-2.0 | 20128 |
| Number_of_word_pairs_correctly_associated-2.0 | 20197 |
| Number_of_puzzles_correct-2.0 | 6373 |
| Number_of_symbol_digit_matches_attempted-2.0 | 23323 |
| Number_of_symbol_digit_matches_made_correctly-2.0 | 23324 |
| Hand_grip_strength_mean_lr-2.0* | 46, 47 |

*Average of Hand_grip_strength_l-2.0 and Hand_grip_strength_r-2.0

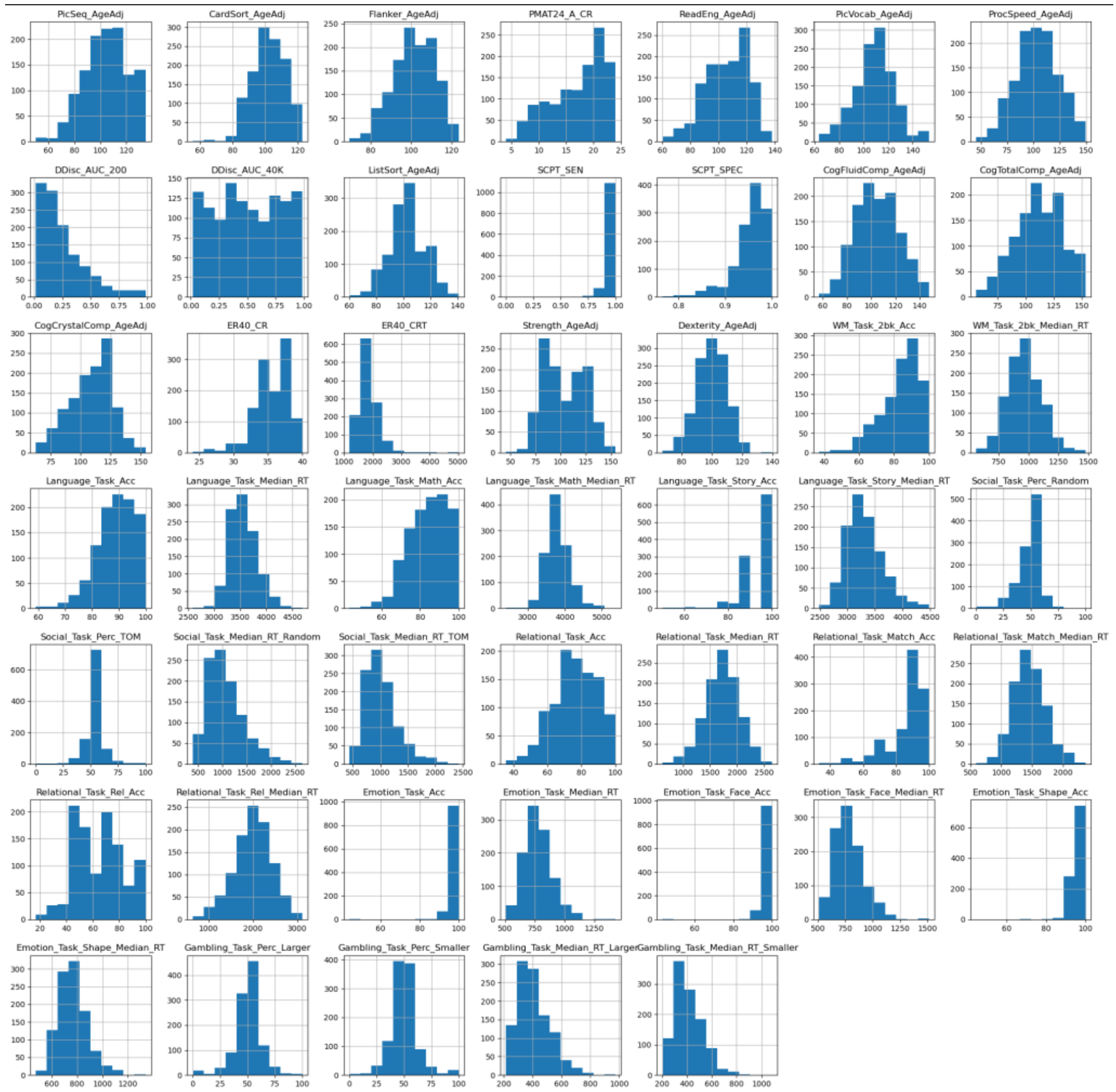


**Supplementary Figure 2.** Distribution of all predicted behaviours in UKB

### 3. Behaviours selected for prediction in ABCD

**Supplementary Table 3**

*Predicted behaviours in ABCD*

| Behaviour |
|---|
| nihtbx_picvocab_fc |
| nihtbx_flanker_fc |

nihtbx_pattern_fc

nihtbx_picture_fc

nihtbx_reading_fc

nihtbx_cryst_fc

pea_ravlt_sd_trial_vi_tc

pea_ravlt_ld_trial_vii_tc

lmt_scr_perc_correct

lmt_scr_perc_wrong

lmt_scr_avg_rt

lmt_scr_rt_correct

tfmri_mid_all_beh_srwpfb_nt

tfmri_mid_all_beh_lrwpfb_mrt

tfmri_mid_all_beh_srwpfb_mrt

tfmri_mid_all_beh_lrwpfb_nt

tfmri_sst_all_beh_total_mssrt

tfmri_nb_all_beh_c2b_rate

tfmri_nb_all_beh_c2b_mrt

tfmri_nb_all_beh_c0b_rate

tfmri_nb_all_beh_c0b_mrt

tfmri_rec_all_beh_place_dp

tfmri_rec_all_beh_negf_dp

tfmri_rec_all_beh_neutf_dp

tfmri_rec_all_beh_posf_dpr

### 4. Excluded fields for UKB sample

Subjects with a history of neurological disease, as reported in (Kweon et al., 2022) and additionally subjects with sleep apnoea were excluded.

Excluded ICD codes:

'G473', 'F00', 'F01', 'F02', 'F03', 'G30', 'G20',
'G21', 'G23', 'G31', 'G32', 'G610', 'G35', 'G37',
'I63', 'G463', 'G464', 'I64', 'I694', 'C70', 'C71',
'D33', 'I60', 'I61', 'I62', 'I691', 'I692', 'I693',
'G060', 'G07', 'I671', 'Q282', 'Q283', 'G80', 'A521',
'A504', 'I64', 'A83', 'A86', 'B011', 'B020', 'B262',
'A85', 'B004', 'B582', 'A84', 'B050', 'B941', 'G04',
'A321', 'G05', 'G40', 'F803', 'S07', 'T040', 'A80',
'A81', 'A82', 'A83', 'A84', 'A85', 'A86', 'A87', 'A88',
'A89', 'G45', 'C70', 'C793', 'D32', 'D33', 'G03', 'A170',
'A171', 'A203', 'G01', 'G02', 'G00', 'G07', 'G122', 'Q05',
'Q760', 'P100', 'I60', 'S066', 'P103', 'G45',
'F'

Self-reported illness code (Data-Coding 6):

1123, 1263, 1262, 1258, 1256,
1261, 1397, 1081, 1032, 1491,
1245, 1425, 1433, 1246, 1264,
1266, 1244, 1583, 1031, 1659,
1247, 1259, 1240, 1524, 1083,
1086, 1082

### 5. Sample size for predicted behaviours in HCP-A

**Supplementary Table 4**

*Description of sample for behavioural prediction in HCP A*

| Behaviour | Sample (Female) | Ages |
|---|---|---|
| Age (in months) | 647 (351) | 36-86 |
| Total cognition (age adjusted) | 550 (308) | 36-86 |
| Crystalised cognition (age adjusted) | 549 (308) | 36-86 |
| Hand grip strength (dominant hand) | 551 (306) | 36-86 |

### 6. Acquisition and preprocessing parameters

**Supplementary Table 5**

*Acquisition and preprocessing of datasets*

| Parameter | Dataset | | |
|---|---|---|---|
| | HCP-A | HCP-YA | UKB |
| Scanner | Siemens Prisma 3T | Siemens ‘Connectom Skyra’ 3T | Siemens Skyra 3T |
| Resting-state sessions | 4 runs across 2 days with AP and PA phase encoding on each day | 4 runs across 2 days with LR and RL phase encoding on each day | a single run acquired in AP direction |
| acquisition time | 488 frames per run (26 min total) | 1200 frames per run (58 min total) | 490 volumes (6 min) |
| TR/TE | 800/37 ms | 720/33 ms | 735/39 ms |
| Acquisition sites | 4 | 1 | 4 |
| gradient distortion correction | yes | yes | yes |
| intensity normalisation | yes | yes | yes |
| motion correction | yes | yes | yes |
| normalisation to MNI | yes | yes | yes |
| artefact removal | ICA-FIX | ICA-FIX | ICA-FIX |
| temporal filtering | bandpass filtered at 0.01 – 0.1 Hz | bandpass filtered at 0.01 – 0.1 Hz | highpass filtering |
| denoising | WM+CSF+GS regression | WM+CSF+GS regression | no |

*Abbreviations; AP: anterior-to-posterior; PA: posterior-to-anterior; LR: left-to-right; RL right-to-left all refer to phase encoding directions*

### 7. Comparison of accuracy to computation time

Given the large number of predictions that were necessary to conduct for our simulation analyses, we first tested algorithms commonly used in the literature (Supplementary Figure 3) to find the best computation time to accuracy trade-off (measured with R2). This was not meant as an exhaustive comparison to identify the perfect pipeline. Four algorithms were tested: linear ridge regression, kernel ridge regression and two flavours of support vector regression all implemented in the Scikit learn library [version 0.24.2, (Pedregosa et al., 2011)]. A 10-fold cross-validation scheme was used to evaluate the performance of all models. Hyperparameter optimization of the alpha regularisation parameter for ridge regression and kernel ridge regression were performed using a nested cross-validation scheme (5-fold cross-validation for kernel ridge and leave-one-out cross-validation for ridge regression) embedded with the 10-fold cross-validation. Next, two implementations of linear support vector regression were tested (sklearn.svm.LinearSVR with squared epsilon-insensitive (L2) loss function and sklearn.svm.SVR; see https://github.com/MartinGell/Prediction_Reliability for details). A heuristic was used to efficiently calculate the hyperparameter C (Helleputte, Paul, & Gramme, 2021): $c = \frac{1}{\frac{1}{n}\sum_{i=1}^{n}\sqrt{G[i,i]}}$ where G = matrix multiplication of features and transpose of features (here functional connectivity).

Prior to training, subjects with behavioural data over +-3SD were removed and neuroimaging features were z-scored within participants (average connectomes from HCP dataset were first transformed back to r values from Fisher-z scores) to keep features consistent across algorithms as this step was required for kernel ridge regression. Within each training fold, neuroimaging features were z-scored across participants before models were trained (using Sklearn’s `pipeline`).

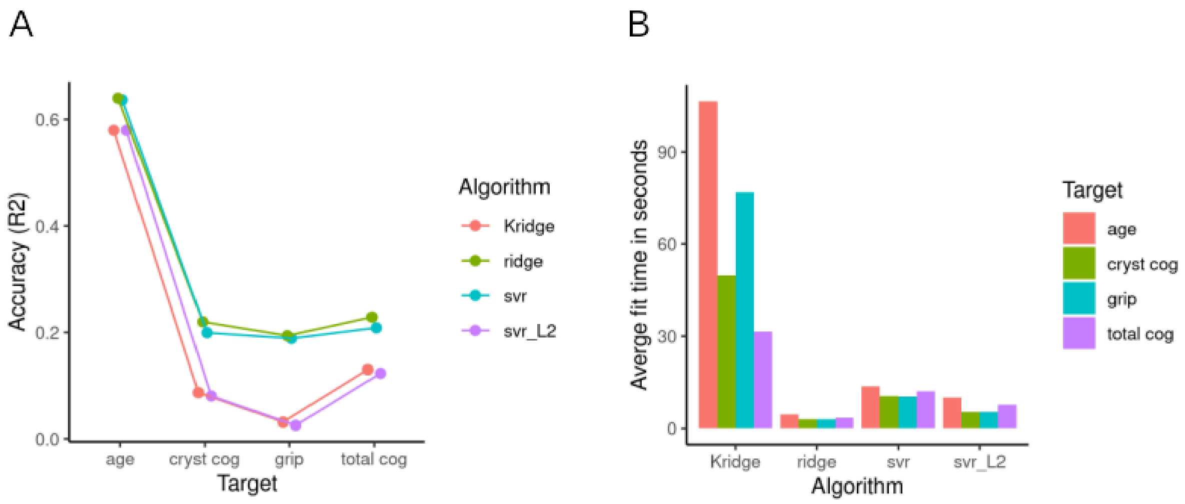


**Supplementary Figure 3. Comparison of algorithms.** (A) Displays prediction accuracy (R2) for all behaviours of interest predicted using different algorithms: Kridge = Kernel ridge regression, ridge = Linear ridge regression, svr = Support vector regression and svr_L2 = support vector regression. (B) Displays average training time of a single mode across cross-validation for all tested behaviours and algorithms.
